# Human co-transcriptional splicing kinetics and coordination revealed by direct nascent RNA sequencing

**DOI:** 10.1101/611020

**Authors:** Heather L. Drexler, Karine Choquet, L. Stirling Churchman

**Affiliations:** Department of Genetics, Blavatnik Institute, Harvard Medical School, Boston, MA 02115

## Abstract

Human genes have numerous exons that are differentially spliced within pre-mRNA. Understanding how multiple splicing events are coordinated across nascent transcripts requires quantitative analyses of transient RNA processing events in living cells. We developed nanopore analysis of CO-transcriptional Processing (nano-COP), in which nascent RNAs are directly sequenced through nanopores, exposing the dynamics and patterns of RNA splicing without biases introduced by amplification. nano-COP showed that in both human and *Drosophila* cells, co-transcriptional splicing occurs after RNA polymerase II transcribes several kilobases of pre-mRNA, suggesting that metazoan splicing transpires distally from the transcription machinery. Inhibition of the branch-site recognition complex SF3B globally abolished co-transcriptional splicing in both species. Our findings revealed that splicing order does not strictly follow the order of transcription and is influenced by cis-regulatory elements. In human cells, introns with delayed splicing frequently neighbor alternative exons and are associated with RNA-binding factors. Moreover, neighboring introns in human cells tend to be spliced concurrently, implying that splicing occurs cooperatively. Thus, nano-COP unveils the organizational complexity of metazoan RNA processing.

## Introduction

Most human genes contain many long introns with degenerate sequence information at splice sites, requiring sophisticated mechanisms to correctly localize and coordinate the excision of multiple introns within the same nascent transcript. This process requires the multi-megadalton spliceosome complex, which assembles in a stepwise fashion at each intron splice site (SS) within a pre-mRNA (Fica and Nagai, 2017; Wahl et al., 2009). Intron splicing comprises two catalytic transesterification steps, which involve assembly and rearrangements of the spliceosome. First, the branch point adenosine attacks the 5’SS, forming an intron lariat. Second, the free end of the 5’SS attacks the downstream 3’SS, joining the exons and releasing the intron lariat.

*In vitro* studies have demonstrated that recruitment and assembly of the spliceosome depends on the predominant gene architecture of an organism. Splicing of short introns (<250 nt) typically occurs through an intron definition model, in which spliceosome factors are initially recruited to the 5’ and 3’ splice sites across an intron to facilitate its splicing. By contrast, long introns (>250 nt) are spliced through an exon definition model, in which splicing factors are recruited to the junctions of an exon at the 5’SS of its downstream intron and the 3’SS of its upstream intron before cross-intron interactions are made to facilitate intron splicing (Berget, 1995; Schneider et al., 2010). Exon definition splicing is presumed to dominate in human cells because most genes contain long introns separated by short exons, whereas in *Drosophila melanogaster*, which has an abundance of short introns, intron definition splicing is more prevalent (Fox-Walsh et al., 2005; Guo et al., 1993).

Much of our knowledge of splicing mechanisms has been gleaned from *in vitro* experiments using externally synthesized pre-mRNA (Black et al., 1985; Braun et al., 2018; Hoskins et al., 2011; Krainer et al., 1984). However, because splicing is predominantly co-transcriptional (Beyer and Osheim, 1988; Khodor et al., 2011; Pandya-Jones and Black, 2009; Wuarin and Schibler, 1994), such studies cannot capture a complete view of *in vivo* splicing. For instance, mutations in the C-terminal domain and trigger loop of RNA polymerase II (Pol II) alter splicing outcomes, suggesting a physical and mechanistic coupling between the transcription and splicing machineries (de la Mata et al., 2003; McCracken et al., 1997). Elucidating the details of the *in vivo* splicing reaction requires analysis of newly synthesized RNAs, which must be carefully purified. A variety of approaches have provided estimates of the length of time between the synthesis of an intron and its removal by splicing in living cells. Measurements of intron synthesis, lariat degradation, and the completion of the splicing reaction by quantitative PCR (Singh and Padgett, 2009), metabolic labeling (Rabani et al., 2014; Windhager et al., 2012), or single-molecule imaging (Coulon et al., 2014; Martin et al., 2013) have revealed that intron removal can take between 30 seconds and an hour in mammalian cells. Although enlightening, these strategies fail to uncover the physical link between transcription and splicing, as well as the patterns of splicing across nascent transcripts. Recent work in the yeast species *S. cerevisiae* and *S. pombe* suggests that splicing is completed nearly immediately after an intron is synthesized (Carrillo Oesterreich et al., 2016; Herzel et al., 2018), but it remains to be seen whether this is also the case in organisms with more complex gene structures and abundant alternative splicing, such as *Drosophila* and humans.

If transcription and splicing are coupled, then the order in which introns are transcribed has been reasoned to influence the order in which they are spliced. In fact, the “first-come, first-served” model of splicing proposes that the first introns transcribed are the first to be committed for splicing (Aebi and Weissman, 1987). However, even though the completion of intron splicing frequently follows a defined order within a gene, the order of splicing is not always concordant with the direction of transcription (Kessler et al., 1993; Kim et al., 2017). Splicing order has the potential to strongly influence alternative splicing decisions (Takahara et al., 2002). For example, whether the splicing of each intron is controlled by a defined kinetic window or instead depends on the splicing status of nearby introns has profound consequences for models of splicing regulation. Control of the order of intron splicing and the coordination of splicing patterns across multi-intron genes are largely unexplored topics that are critical to our understanding of alternative splicing.

Current gaps in our understanding of co-transcriptional splicing regulation are mainly due to three challenges in the quantitative analysis of nascent transcripts. First, nascent RNA is a very small fraction (<0.5%) of cellular RNA, presenting a purification challenge that can lead to overestimates of co-transcriptional splicing measures if samples are contaminated with spliced mature RNA. Second, length biases that arise during the enzymatic steps of library preparation can systematically distort relative measurements of spliced and unspliced RNAs that differ in size by the length of an intron. Third, traditional short-read sequencing does not provide isoform information across RNA molecules, limiting its ability to detect multi-intron splicing patterns.

Here, we describe nanopore analysis of CO-transcriptional Processing (nano-COP), a technique designed to directly probe the dynamics and regulation of pre-mRNA splicing *in vivo*. nano-COP uses direct RNA nanopore sequencing to determine the native isoform of long nascent RNA molecules without amplification-associated biases. Application of nano-COP to human and *Drosophila* cells revealed global features of RNA processing in both species. We observed striking differences in splicing kinetics between the two species suggesting that intron definition splicing is more rapid than exon definition splicing. We also discovered global patterns in the order of intron splicing and identified coordinated patterns of splicing in human cells. Thus, nano-COP represents an effective strategy for exposing the critical molecular processes that occur during transcription in living metazoan cells.

## Results

### Unveiling the nascent transcriptome with direct RNA nanopore sequencing

We developed nanopore analysis of CO-transcriptional Processing (nano-COP) in order to observe nascent RNA isoforms as they are transcribed. In this method, stringent purification of nascent RNA followed by long-read nanopore sequencing connects intron splicing to Pol II positions at the 3’ ends of reads, revealing the relationship between transcription and splicing in living cells (Figure 1A). To sequence long RNA molecules without amplification-associated biases, we adopted a direct RNA sequencing approach using Oxford Nanopore Technologies’ MinION instrument (Garalde et al., 2018). Because nanopore sequencing is lower-throughput than short-read sequencing, and even low background levels of mature spliced RNA can result in overestimation of co-transcriptional splicing measures, we developed a stringent nascent RNA purification strategy by combining two complementary techniques: cellular fractionation (Pandya-Jones and Black, 2009; Wuarin and Schibler, 1994) (Figure S1A) and 4-thiouridine (4sU) pulse labeling (Dölken et al., 2008; Schwalb et al., 2016).

**Figure 1.**
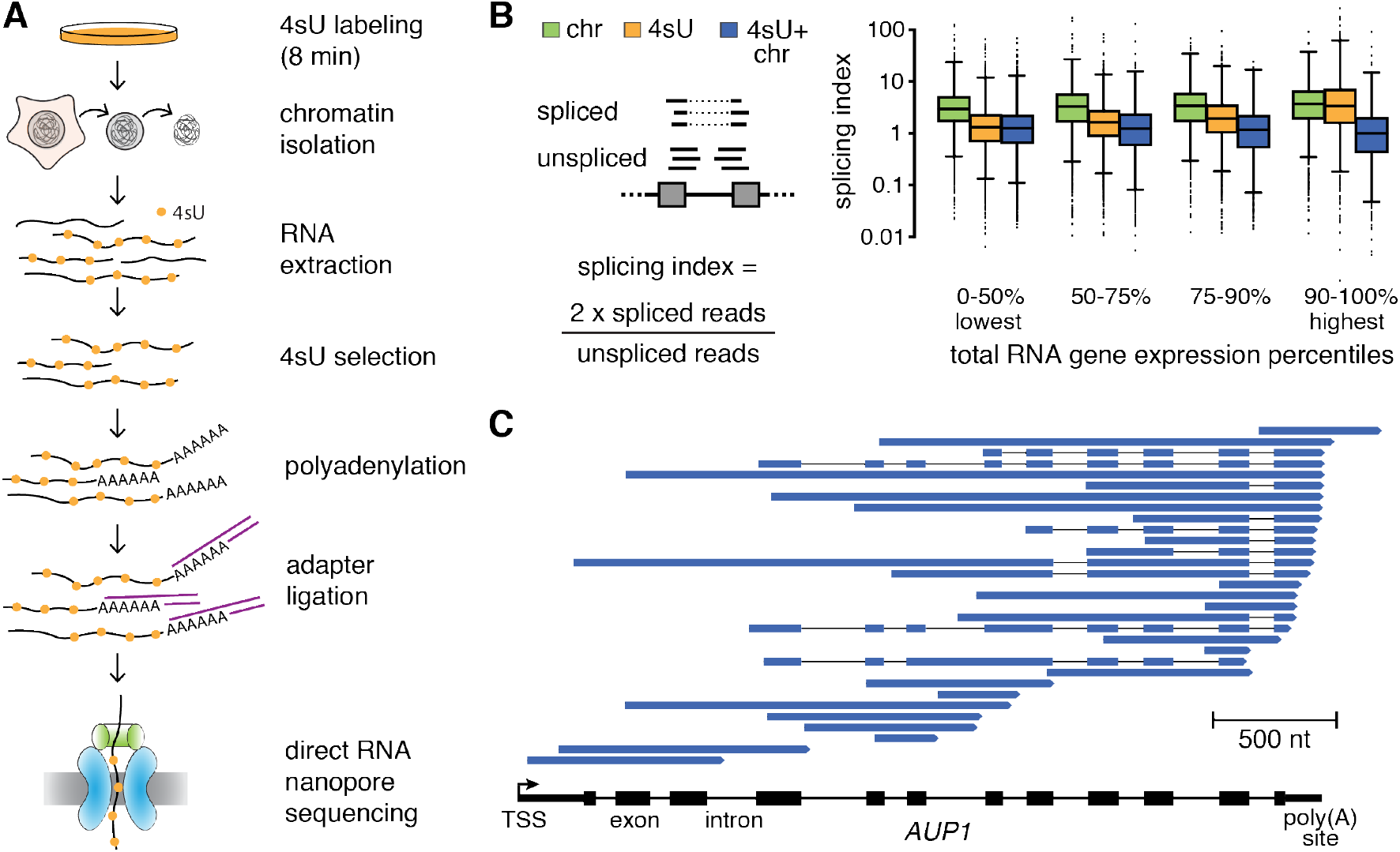
Capturing the nascent transcriptome in human cells with nano-COP. (A) Schematic of the nano-COP protocol with nascent RNA purification and direct RNA nanopore sequencing. Cells are labeled with 4-thiouridine (4sU); cellular fractionation is performed to isolate chromatin-associated RNA; and 4sU labeled chromatin-associated RNA is selected through biotinylation and affinity purification. Subsequently, the purified RNA is polyadenylated using *E. coli* poly(A) polymerase. The sample is subjected to direct RNA library preparation, including 3’ end adapter ligation, followed by nanopore sequencing. (B) Splicing index, which represents the proportion of spliced transcripts in Illumina sequencing datasets, is plotted within different percentiles of total RNA gene expression. The distribution of 4sU labeled chromatin-associated RNA (4sU+chr) differs significantly (t-test p-value < 0.05) between chromatin-associated RNA (chr) and 4sU labeled RNA (4sU) at all gene expression levels. Gene expression percentiles are based on total RNA expression levels out of 9659 genes that have at least 25 reads spanning splice junctions in all datasets. (C) Representative nano-COP reads aligned to the *AUP1* gene in human K562 cells. The gene structure is represented from the transcription start site (TSS) to the poly(A) site, with black boxes representing exons and lines representing introns. Within the reads, blue boxes represent read coverage and black lines represent skipped coverage due to splicing.

After cells are labeled with 4sU for 8 minutes, they are lysed and subjected to sequential centrifugation steps that yield the chromatin, nucleoplasm and cytoplasm cellular fractions. RNA purified from the chromatin fraction is then subjected to 4sU purification. Even for much longer periods than 8 minutes, 4sU treatment does not show a detectable impact on splicing (Schofield et al., 2018) (Figure S1B–C), supporting the use of this metabolic labeling approach to quantify splicing rates (Pai et al., 2017; Rabani et al., 2014). The combination of cellular fractionation and 4sU purification enriches for RNAs that are both recently transcribed and localized to the site of transcription. Moreover, the use of both methods in tandem yields higher amounts of unspliced RNA at all gene expression levels compared to either method alone, resulting in a substantial improvement in the purification of nascent RNA (Figures 1B and S1D–E).

During direct RNA nanopore sequencing, each RNA molecule is recruited through the nanopore via an adapter ligated to its polyadenylated [poly(A)] tail, and is consequently read from 3’ to 5’ (Garalde et al., 2018). Because we expect most 4sU-labeled chromatin-associated RNAs to be nascent, and therefore not yet polyadenylated, we add poly(A) tails enzymatically using *E. coli* poly(A) polymerase before ligating the adapter (Figure 1A). After sequencing and aligning reads to the genome, we can distinguish RNAs that have completed transcription with endogenous poly(A) tails from nascent RNAs with artificial poly(A) tails based on where the 3’ end of the RNA aligns.

We first applied nano-COP to human chronic myeloid leukemia K562 cells. Analysis of datasets from biological triplicates confirmed that gene expression levels were reproducible across replicates (Pearson’s R = 0.81; Figure S2A) and correlated with short-read RNA sequencing coverage by the Illumina platform (Pearson’s R = 0.79; Figure S2B) even though nanopore sequencing is lower throughput (Figure S2C). Read accuracies were nearly identical to those of nanopore datasets acquired using poly(A)-selected mature RNA (Workman et al., 2018), indicating that 4sU does not have a detectable effect on base calling (Figure S2D-E). We obtained a wide range of read lengths (median, 671 nt; longest, 7420 nt; Figure S2F-G), comparable to previously reported direct RNA read lengths from mRNA (Workman et al., 2018). Over 25% of nascent RNA nanopore reads were greater than 1 kb in length, enabling detection of co-transcriptional RNA processing over long ranges. To assess the variation in processing dynamics between species with diverse gene architectures and splicing regulation, we also applied nano-COP to *D. melanogaster* S2 cells (Figure S3A).

Each nano-COP sequencing run yielded ∼100,000 mapped reads that aligned to >11,000 genes, with a median of one read mapping per gene (Figure S2C,H); highly expressed genes with many read alignments showcase the typical diversity of nascent transcript isoforms in human K562 cells (Figure 1C). The positioning of read 3’ ends within genes, along with the presence of unspliced introns, indicate that the sequenced reads originated from RNAs that were in the midst of being transcribed and processed.

In addition to canonical coding genes, nano-COP exposes the transcription and RNA processing of noncoding RNAs. For instance, we observed nano-COP reads that start and end at the boundaries of precursor micro RNAs (pre-miRNA) within unspliced introns (Figure S3B), supporting previous findings that pre-miRNA cleavage can occur before intron splicing (Kim and Kim, 2007). We also detected cases of antisense transcription across long noncoding RNAs (Figure S3C). Although current nanopore sequencing technology does not provide sufficient depth for analysis of single genes or introns, in the aggregate, nano-COP data has the potential to yield insights into fundamental features of global splicing dynamics.

### Pol II mapping with nano-COP

In native elongating transcript sequencing (NET-seq)-based approaches, sequencing of the 3’ ends of nascent RNAs enables high-resolution mapping of Pol II genome-wide (Churchman and Weissman, 2011; Mayer et al., 2015; Nojima et al., 2015). To determine whether nano-COP also measures Pol II position, we sought to verify that the start of nano-COP reads represents the 3’ ends of nascent RNA. We find thattranscript 3’ ends aligned mostly within gene bodies (Figure S4A-B), signifying that they originated mainly from RNAs that were still undergoing transcription. By contrast, libraries constructed from 4sU-labeled chromatin-associated RNA without enzymatic addition of a poly(A) tail aligned predominantly to annotated poly(A) sites (Figure S4C), and thus represented pre-mRNA that remained associated with chromatin after transcription is complete (Brody et al., 2011). Furthermore, the poly(A) tails at the start of each read were shorter when aligned within gene bodies and longer when aligned to poly(A) sites, demonstrating that the former were enzymatically polyadenylated and therefore distinguishable from the naturally polyadenylated RNAs (Figure S4D) (Loman et al., 2015; Workman et al., 2018). Importantly, despite the high error rates of direct RNA nanopore sequencing, especially on homopolymers like poly(A) stretches, most 3’ ends of synthetic RNAs with encoded poly(A) tails correctly mapped within 25 nt after sequencing (Figure S4E) (Garalde et al., 2018). Based on these results, we conclude that the 3’ ends of nascent transcripts with artificial poly(A) tails largely represent active transcription sites.

In addition to Pol II position, nano-COP also captures intermediates of the splicing reaction (i.e. free ends of upstream exons after the first catalytic step of splicing), providing insights into the regulation and efficiencies of the catalytic steps of splicing (Burke et al., 2018; Chen et al., 2018) (Figure S4F-G). The alignment positions of nascent RNA 3’ ends allow for the computational isolation of reads arising from RNAs that are being actively transcribed from those that have completed transcription (i.e., RNA 3’ ends aligning to poly(A) sites) or have undetermined transcription status (i.e., RNA 3’ ends aligning to splice sites).

### nano-COP reveals the relationship between transcription and splicing

With long read lengths, nano-COP can simultaneously determine the position of Pol II and the splicing status of each intron in a nascent RNA. To determine the physical proximity between splicing catalysis and transcription, we tallied the splicing status of introns within each read as a function of Pol II position (i.e., the 3’ end of nascent RNA) (Figure 2A). Focusing initially on constitutively spliced introns, we found that in human K562 cells, Pol II transcribed several kilobases (kb) before introns were spliced (Figure 2B and S5A–C). nano-COP analysis in *Drosophila* S2 cells revealed that splicing of most introns occurred more proximally to Pol II, such that ∼25% percent of introns were spliced within 1 kb, and more than 50% were spliced within 2 kb from the intron 3’SS (Figure 2B and S5A–B,D). These results did not change when the first and last introns of all genes were removed from the analysis (Figure S5E).

**Figure 2.**
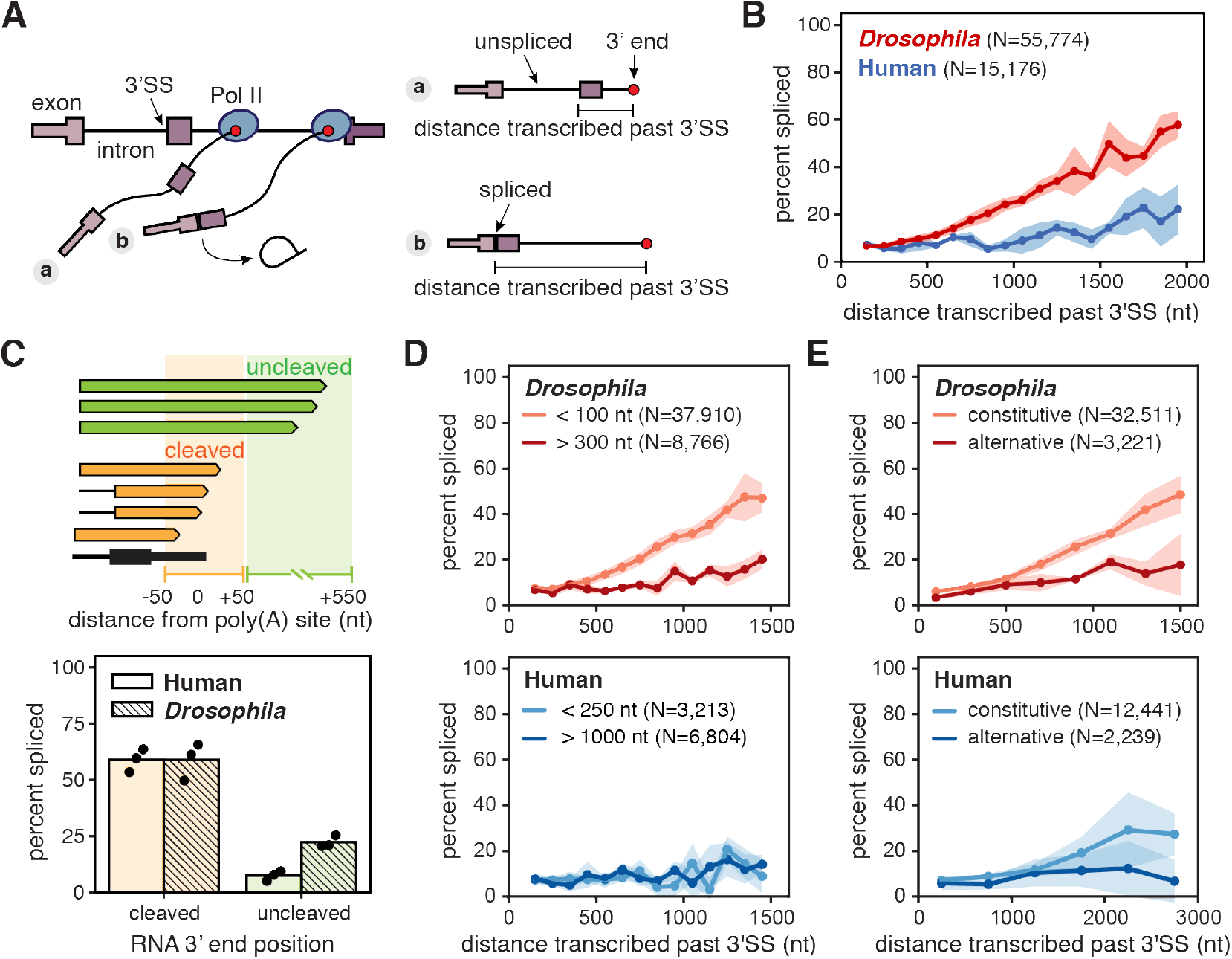
nano-COP measures the physical proximity between transcription and splicing. (A) Cartoon depicting measurements of distance transcribed past the 3’ splice site (3’SS) and splicing status using nano-COP data from two nascent transcripts. (B) Global analysis of distance transcribed from the 3’SS and the percent of spliced molecules in human K562 (blue) and *Drosophila* S2 (red) cells. (C) Cartoon (top) represents example reads with 3’ ends near gene ends and the regions that differentiate “cleaved” (orange, 50 nt upstream and downstream of poly(A) sites) from “uncleaved” (green, 50-500 nt downstream of poly(A) sites) transcripts. Bar plot (bottom) depicts the proportion of reads with spliced terminal introns in the “cleaved” and “uncleaved” pools of transcripts for human K562 (solid) (N=10,669 cleaved; N=1,068 uncleaved) and *Drosophila* S2 (hashed) (N=9,534 cleaved; N=6,062 uncleaved) cells. Black points represent results from three biological replicates. (D) Global measurement of distance transcribed before splicing as a function of intron size. (E) Global analysis of transcribed distance from 3’SS and the percent of spliced molecules separated by alternative splicing status of neighboring exons. Introns that do not neighbor an alternatively spliced exon are labeled as ‘constitutive’ while introns that do neighbor an alternatively spliced exon are labeled as ‘alternative’. Numbers (N) in B, D, and E represent the number of introns within reads that were used to calculate the distance transcribed before splicing plots. Shaded regions (in B, D, and E) represent standard deviation across three biological replicates.

To assess the extent of post-transcriptional splicing in both species, we focused on the relationship between terminal intron splicing and transcription termination. In human cells, terminal introns were almost exclusively spliced post-transcriptionally (Figure 2C). This finding corroborates established functional links between polyadenylation and terminal intron splicing in vertebrates (Berget, 1995; Niwa and Berget, 1991). In *Drosophila* cells, ∼25% of terminal intron splicing occurred prior to cleavage and polyadenylation (Figure 2C), consistent with the faster co-transcriptional splicing in those cells.

Variations in splicing kinetics between species are likely due to mechanistic differences in splicing regulation. In *Drosophila* cells, short introns (<100 nt) were excised faster than long introns (>300 nt) (Figure 2D and S5F) (Khodor et al., 2011; Pai et al., 2017), and introns neighboring alternative exons were spliced more slowly than those neighboring constitutive exons (Figure 2E). However, this difference was partly due to the tendency of long introns to flank alternative exons in *Drosophila* (Figure S5G) (Fox-Walsh et al., 2005). The abundance of short introns that are rapidly and constitutively spliced in *Drosophila* cells implies that splicing through intron definition is more efficient than exon definition. By contrast, we did not observe the same length dependence for splicing kinetics in human cells (Figure 2D), and only a slight difference (if any) between constitutive and alternative introns (Figure 2E). A difference at greater distances of Pol II transcription may be found once increased read lengths permit them to be resolved. Nonetheless, these data indicate that the existence of other modes of splicing regulation in humans are likely to be responsible for the differences in splicing kinetics relative to *Drosophila*.

### SF3B activity is required for co-transcriptional splicing

Splicing factors are commonly mutated in certain cancer types (Quesada et al., 2011; Wang et al., 2011; Yoshida et al., 2011), motivating the characterization of splicing inhibitor compounds that exert antitumor activity. For example, mutations in SF3B1, a member of the SF3B complex that performs branch point recognition prior to the first catalytic step of splicing, alter splicing activity and are sufficient to cause cancer phenotypes in mice (Obeng et al., 2016). Several SF3B1 inhibitors, both naturally occurring and synthetically derived, have antitumor properties (Bonnal et al., 2012). However, even though this protein plays a critical role in all splicing reactions, inhibitors of SF3B1 change alternative splicing rather than global intron retention in steady state pools of RNA (Teng et al., 2017; Vigevani et al., 2017). It remains unclear whether this response is due to differential impacts of SF3B1 on splicing or variation in RNA turnover or surveillance processes.

To explore this issue, we investigated the impact of the small molecule pladienolide B (PlaB), which competitively binds to SF3B1 (Cretu et al., 2018; Kotake et al., 2007). Consistent with previous studies of SF3B1 inhibitors, RT-PCR analysis revealed that PlaB treatment caused no detectable changes in splice isoforms of cytosolic transcripts (Figure 3A). However, when we purified and analyzed chromatin-associated RNAs after PlaB treatment, we observed an accumulation of unspliced transcripts (Figure 3A). We analyzed published datasets of RNAs that are closer to the site of transcription [nucleoplasm RNA-seq (NP-seq) and mammalian NET-seq (mNET-seq)] and observed only a modest decrease in the percent of spliced reads after 4 hours of treatment with PlaB (Figure 3B-C) (Nojima et al., 2015). These results indicate that either PlaB does not have a global influence on splicing or that challenges in analyzing nascent RNA conceal its impact.

**Figure 3.**
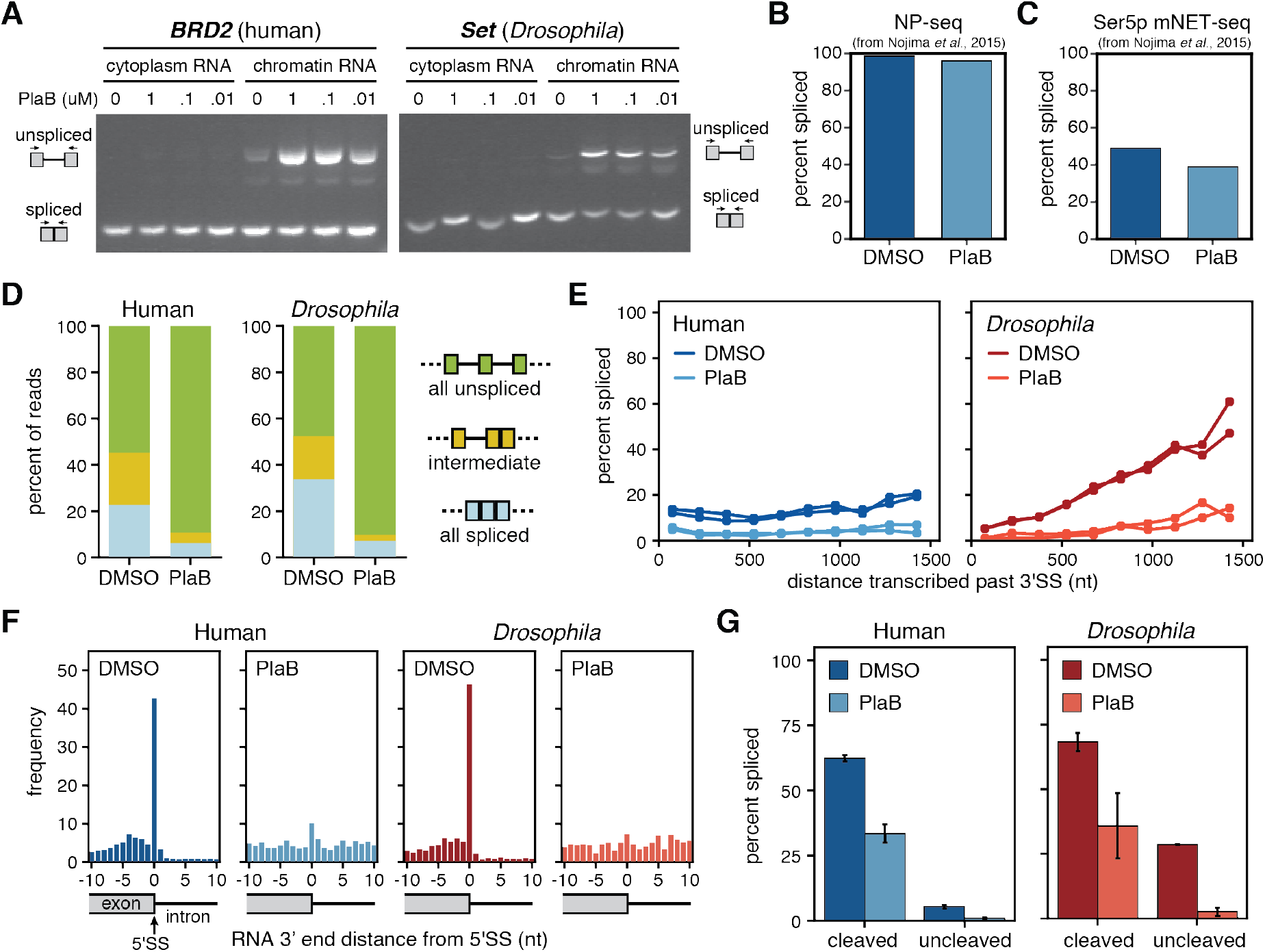
Co-transcriptional splicing is abolished with the splicing inhibitor PlaB. (A) RT-PCR of example genes in cytoplasmic and chromatin RNA extracted from human K562 cells (left) and *Drosophila* S2 cells (right) treated with 0.1% DMSO or different concentrations of PlaB (1 uM, 100 nM, and 10 nM). (B) NP-seq and (C) mNET-seq datasets from cells treated with 1 μM PlaB or 0.1% DMSO for 4 hours from (Nojima et al., 2015) were compared for the global percent of spliced reads (spliced reads / total reads aligning to 3’SS junctions). (D) Distribution of splicing patterns found in cells treated with 0.01% DMSO versus 0.1 μM PlaB for 1 hour. In nano-COP reads spanning at least two introns, “all spliced” represents reads where every intron within the read is spliced, “intermediate” represents reads where at least one intron is spliced and one intron is unspliced, and “all unspliced” represents reads where every intron is present and therefore not spliced. (E) Global analysis of distance transcribed past the 3’SS and the percent of spliced molecules in cells treated in biological duplicates with DMSO versus PlaB. (F) The frequency of RNA ends (the 3’ end of the RNA when aligned) around intronic 5’SS represented as histograms in human and *Drosophila* cells treated with DMSO or PlaB. High coverage of nascent RNA 3’ ends at 5’SS likely represents free exon ends between the first and second catalytic steps of the splicing reaction. (G) The proportion of reads with spliced terminal introns in the “cleaved” (50 nt upstream and downstream of poly(A) site) and “uncleaved” (50-500 nt downstream of poly(A) site) pools of transcripts from cells treated with DMSO versus PlaB. Bars represent the range from two biological replicates. In B,C,E-G human K562 samples are represented in blue; *Drosophila* S2 samples in red; DMSO samples as the darker shade; and PlaB samples as the lighter shade.

To directly investigate how splicing kinetics are affected by interference with SF3B, we performed nano-COP and examined the influence of PlaB on co-transcriptional splicing. When both human and *Drosophila* cells were treated with PlaB for 1 hour, the proportion of spliced nano-COP reads decreased globally (∼90% of reads are unspliced; Figure 3D). In PlaB-treated cells from both species, Pol II transcribed > 1.5 kb before there was any evidence of splicing, indicating a near-complete elimination of co-transcriptional splicing (Figure 3E). Furthermore, intermediates between the first and second catalytic steps of the splicing reaction, represented by RNAs with 3’ ends at 5’SSs, were abolished in the presence of the splicing inhibitor (Figure 3F). Finally, in PlaB-treated cells, terminal introns exhibited no evidence of splicing prior to transcript cleavage and polyadenylation (Figure 3G). Thus, PlaB is a potent and global inhibitor of pre-mRNA splicing. These results also illustrate the capacity of nano-COP to observe rapid changes in co-transcriptional splicing dynamics.

### Intron splicing does not strictly follow the order of transcription

To determine the extent to which transcription impacts splicing kinetics, we analyzed reads spanning pairs of introns in which one intron was spliced and the other was not (Figure S6A, “intermediate”). In contrast to the “first-come, first-served” model of splicing (Aebi and Weissman, 1987), inspection of genes with high coverage frequently revealed examples of downstream introns that were transcribed last but spliced first, such as the third intron of human *EIF1* (Figure 4A).

**Figure 4.**
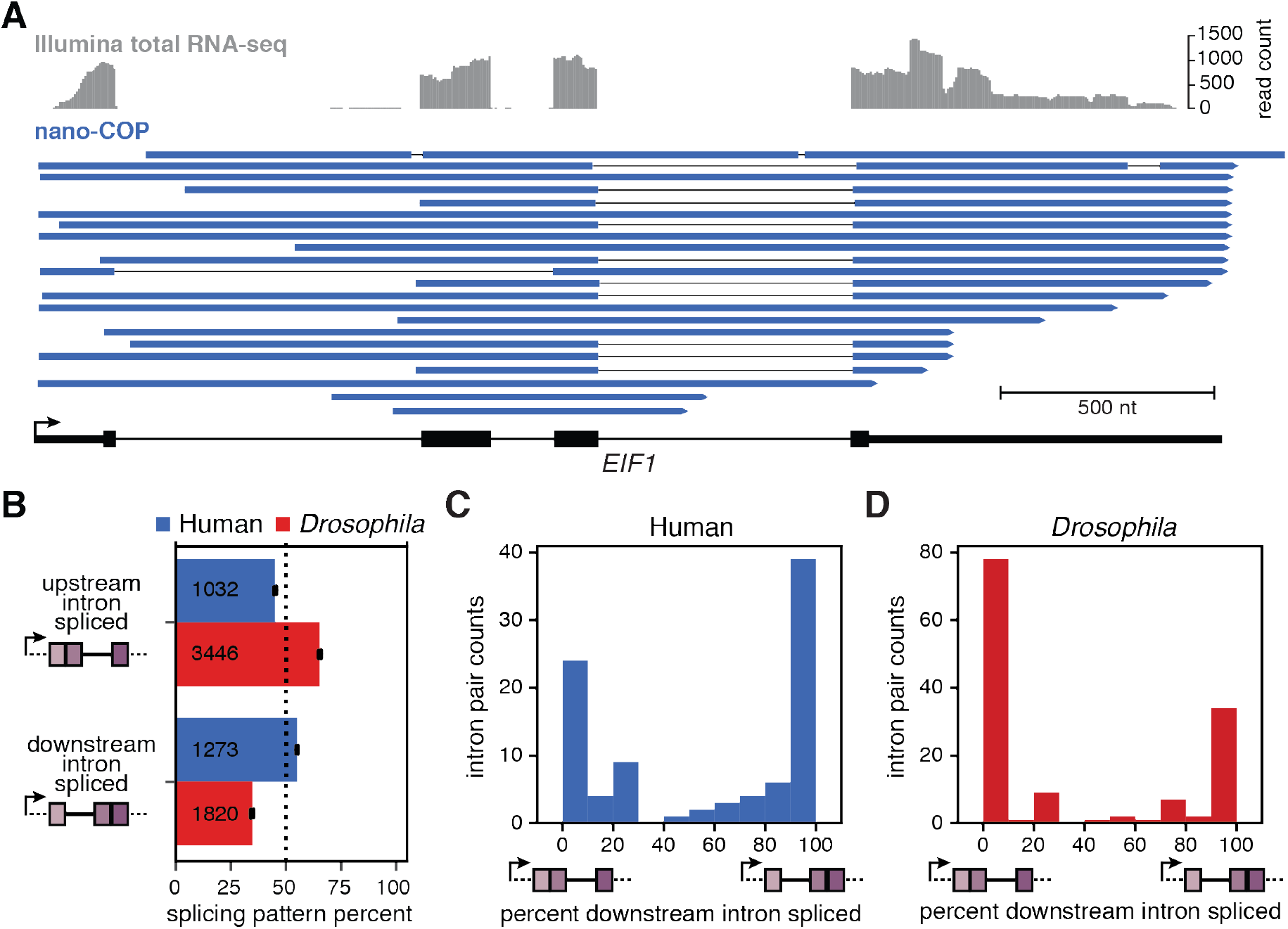
Order of transcription does not strictly dictate splicing order. (A) Illumina total RNA-seq coverage (top, grey) and nano-COP reads (bottom, blue) that align to all three introns in the *EIF1* gene from human K562 cells. (B) Measurements of the order of splicing between neighboring intron pairs in both human (blue) and *Drosophila* (red) cells. For reads that span two or more introns, the top two boxes represent the proportion of reads in which the upstream intron within a pair is spliced first. The bottom two boxes represent the opposite situation, in which the downstream intron within a pair is spliced first. Black bars represent the standard error of the mean across three biological replicates. Values within boxes represent the number of datapoints used to calculate splicing order percentages for each subset. (C–D) Frequency of spliced patterns within intron pairs that have at least 4 reads aligning to both introns in (C) human K562 (N=92 pairs) and (D) *Drosophila* S2 (N=135 pairs) cells.

To measure splicing order globally, we analyzed all reads that spanned intron pairs and found that splicing in human cells is not influenced by transcription order (Figure 4B and S6B), consistent with earlier analyses (Kessler et al., 1993; Kim et al., 2017). In fact, 55% of the time, the downstream intron within a pair was spliced first, indicating that human introns have a slight bias in favor of splicing from 3’ to 5’, in reverse of the transcription order. By contrast, in *Drosophila* cells, the upstream intron tended to be spliced first (65% of the time). This is consistent with the faster co-transcriptional splicing kinetics in this species, as the upstream intron has the possibility of being spliced before the 3’ SS of the downstream intron is transcribed (Figure 4B and S6B). Because the median nanopore read length is less than 1 kb, the introns examined in this analysis tended to be shorter than the natural intron distributions of the two genomes (Figures S2F-G and S6C). However, analysis of splicing intermediates (in which the 3’end of the RNA aligns to a 5’SS) is not similarly constrained by intron length (Figure S6D); for reads corresponding to splicing intermediates, the downstream intron is in the process of being spliced regardless of its length. A large proportion of upstream introns within splicing intermediate reads were not spliced (Figure S6E), further confirming that splicing does not always follow the order of transcription.

We next asked whether splicing patterns are variable or consistent across the same pair of introns, implying that splicing is stochastic or regulated, respectively. Even with the low coverage of nanopore sequencing, we obtained >75 intron pairs in the human and *Drosophila* datasets that were covered by at least four reads exhibiting intermediate splicing patterns. In the vast majority of these intron pairs, every read had the same splicing pattern (Figure 4C-D). Thus, although splicing order does not always follow the direction of transcription, especially in human cells, it does appear to be regulated.

### Intron splicing is influenced by cis and trans regulatory elements

Given that transcription order does not play a major role in splicing order, especially in human cells, we asked whether cis-regulatory features are associated with patterns of intron removal. We found that within pairs, spliced introns tended to be shorter (Figures 5A and S6F) and have stronger splice site scores (Figures 5B-C and S6G-H) than the unspliced introns. These trends were especially pronounced in *Drosophila* cells, but could be observed to a lesser extent in the human dataset.

**Figure 5.**
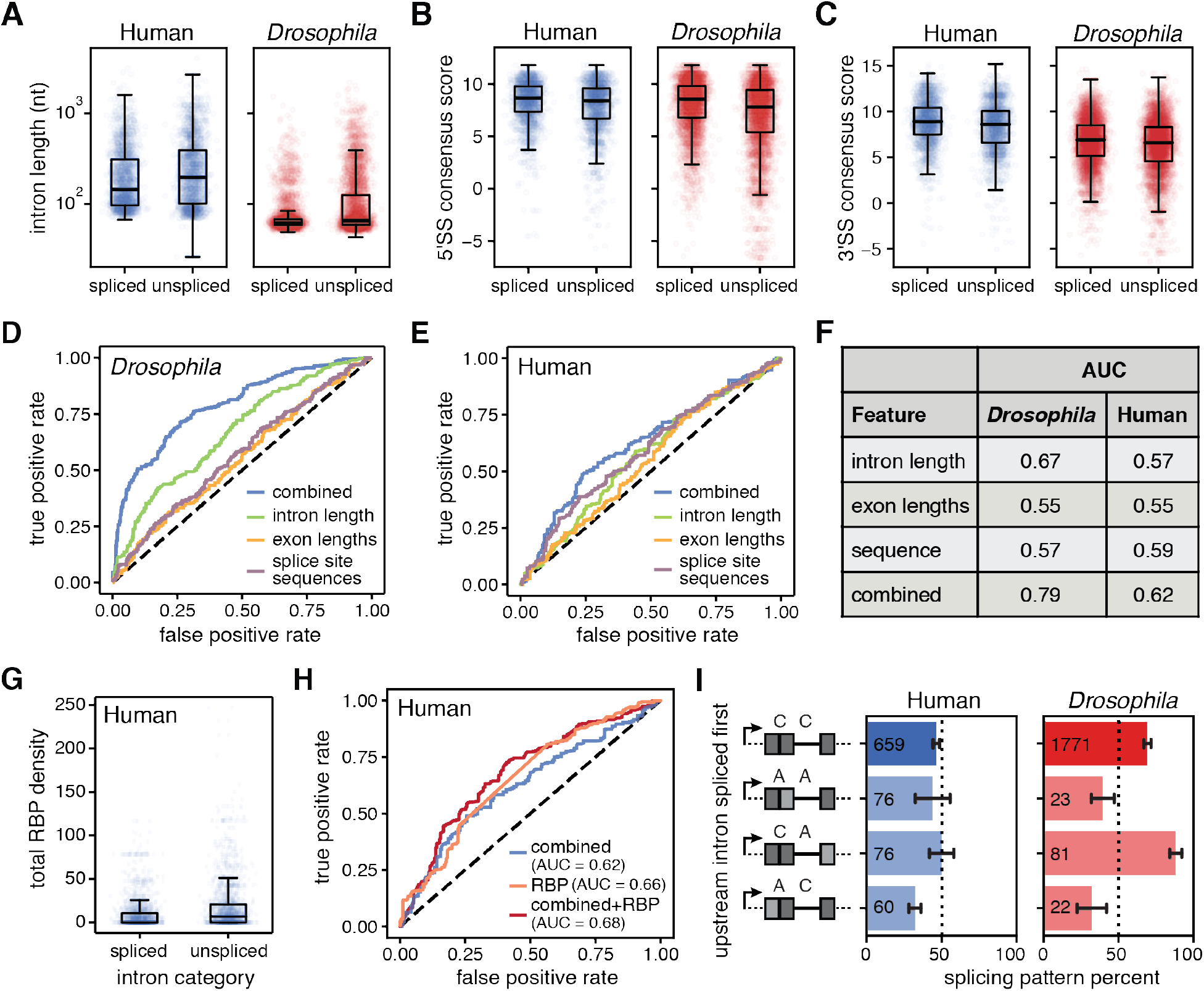
Cis and trans regulatory elements influence splicing order. (A-C) Distributions of (A) intron lengths (human t-test p-value = 3 × 10^−15^; *Drosophila* t-test p-value = 8 × 10^−100^), (B) 5’ splice site (5’SS) consensus sequence scores (human t-test p-value = 5 × 10^−13^; *Drosophila* t-test p-value = 4 ×10^−72^), and (C) 3’SS consensus sequence scores (human t-test p-value = 1 × 10^−13^; *Drosophila* t-test p-value = 3 × 10^−17^) in spliced versus unspliced introns within pairs. (D, E) Receiver Operating Characteristic (ROC) curve from a random forest classifier that measures the predictive value of intron features on splicing order in (D) *Drosophila* S2 cells and (E) human K562 cells. (F) Area under the curve (AUC) measurements for ROC curves from random forest models in D and E. (G) Total RNA binding protein (RBP) density across introns was measured using eCLIP peaks from (Van Nostrand et al., 2018). For all reads that span two or more introns where one is spliced and the other is not spliced, total RBP occupancy was determined as the sum of fold change over input for all peaks from all RBPs in the intron and plotted as a boxplot for the spliced and unspliced introns separately (paired t-test p-value = 1 × 10^−47^). (H) ROC curve from a random forest classifier that measures the predictive value of combined intron features and RBP density on splicing order in human K562 cells. (I) The order of intron splicing across gene regions that undergo alternative splicing. For each splicing pattern, the proportion of cases where the upstream intron is spliced first is represented. Dark gray boxes represent constitutive exons; light gray boxes represent alternative exons; C, intron that neighbors constitutively spliced exons; A, intron that neighbors an alternatively spliced exon. Values within boxes represent the number of datapoints in each category.

To determine whether these global trends strongly influence splicing order, we tested their predictive power using a random forest model. We found that intron lengths, neighboring exon lengths, and splicing consensus sequences were the strongest predictors of splicing order (Supplementary Table 3). Combining these three features within the model improved the prediction, indicating that multiple cis-regulatory features are involved in regulation of splicing patterns (Supplementary Table 3). The combined model for *Drosophila* splicing performed quite well (AUC = 0.79, 75% prediction accuracy) considering how few parameters were included, indicating that cis-regulatory elements play critical roles in splicing order (Figure 5D,F). In human cells, the combined model for splicing did not perform as well (AUC = 0.62, 62% prediction accuracy) (Figure 5E–F), suggesting that additional cis-regulatory elements and/or trans-regulatory factors play more important roles in controlling exon-definition splicing in this species.

To assess the influence of trans-regulatory factors on splicing order in humans, we analyzed a large eCLIP dataset with occupancy measurements for 120 RNA-binding proteins (RBP) in K562 cells (Van Nostrand et al., 2018). By aggregating signal from all RBPs, we found that total occupancy differed significantly between spliced and unspliced introns within pairs (Figure 5G). Examination of individual proteins revealed that 19 RBPs were significantly more likely to be bound to unspliced introns than spliced introns within pairs, both in terms of the number of introns bound and RBP occupancy levels. Nine of these RBPs are members of the core spliceosomal machinery, and three are splicing regulators. The rest are uncharacterized RBPs or factors with known roles in other areas of RNA biology (Supplemental Table 4 and Figure S6I). Combining total RBP occupancy with the above cis-regulatory features improved the prediction of splicing order with the random forest model (Figure 5H), indicating a role for trans factors in regulating splicing order in human cells. One explanation for the higher RBP density in unspliced introns is that introns with delayed splicing are more likely to be present in the eCLIP dataset. However, some of the enriched RBPs in unspliced introns are members of the activated spliceosome (B^act^) (Kastner et al., 2019), suggesting that these introns are prepared for splicing but that something, perhaps another regulatory factor, is delaying the subsequent reactions.

The order of intron splicing can control alternative splicing decisions (Takahara et al., 2002), so we also investigated the splicing order of intron pairs neighboring alternatively spliced exons. Across intron pairs, introns neighboring constitutively spliced exons were frequently spliced before the introns neighboring alternatively spliced exons (Figure 5I). Because the splicing of one intron before another could alter the availability of cis-regulatory elements that influence the splicing regulation in a neighboring exon, this bias has important implications for models of regulated splicing.

### Splicing is coordinated across multi-intron genes in humans

*In vitro* single-molecule studies have demonstrated the capacity for synergistic spliceosomal assembly across human introns (Braun et al., 2018), motivating an analysis of higher-order patterns of splicing *in vivo*. Long-read sequencing enables analysis of splicing patterns of more than two introns. Taking advantage of this capability, we assessed the frequencies of all possible splicing patterns for reads that spanned three or more introns and from RNAs in the process of being spliced (i.e., that contained a mix of spliced and unspliced introns).

In *Drosophila* cells, the splicing of multiple sequential introns tended to follow the direction of transcription (Figure 6A-B), consistent with our analysis of intron pairs (Figure 4B). In human cells, by contrast, intron splicing followed a defined splicing pattern that was not necessarily concordant with the direction of transcription (Figure 6A-B). Furthermore, adjacent introns in human cells were also more likely to have the same splicing status than separated introns, supporting a model that proximal introns are coordinately spliced (Figure 6C). These patterns did not change when the first and last introns of all genes were removed from the analysis (Figure S7A-B). Intriguingly, the patterns of intron splicing in human cells showed that the most upstream and downstream introns in each read were more frequently spliced, suggesting that coordinated splicing begins from the ends of transcripts and works inward. Together, these results reveal that higher-order coordinated patterns of splicing are orchestrated across nascent transcripts in human cells.

**Figure 6.**
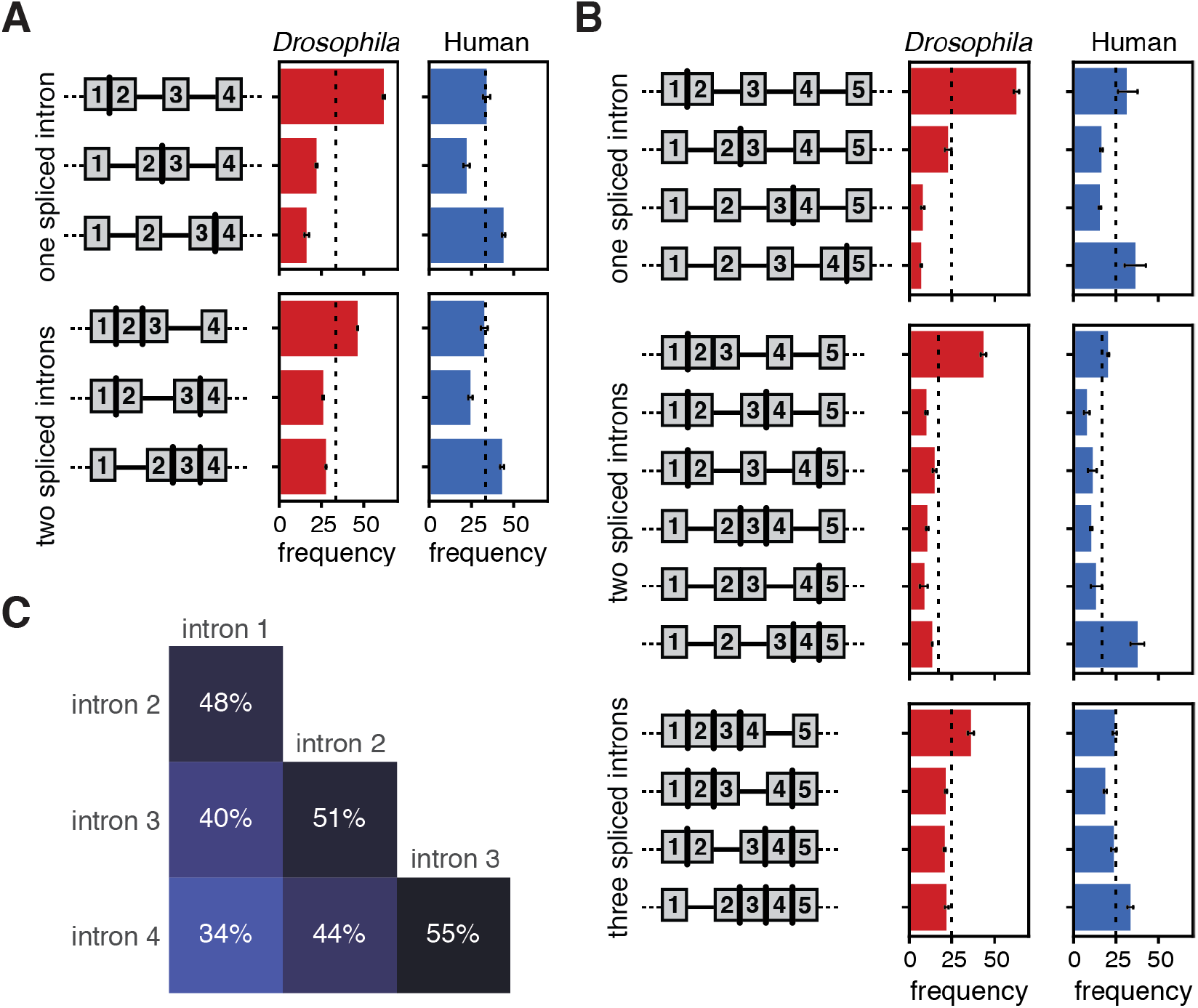
Splicing is coordinated across neighboring introns in human cells. (A) Cartoons of each possible intermediate splicing combination across three sequential introns appear to the left of bar plots representing the percentage of each splicing combination among nano-COP reads from *Drosophila* S2 cells (red, N=2,829) and human K562 cells (blue, N=1,622). The dotted lines at 33% represent the expected percentage if each intron combination were distributed equally. (B) Distribution of splicing patterns in reads that span at least four introns in *Drosophila* S2 cells (red, N=1,214) and human K562 cells (blue, N=956). Dotted lines at 25% for one and three spliced introns and 16.7% for two spliced introns represent the expected percentage if each intron pattern were distributed equally. Error bars represent standard error of the mean across three biological replicates. (C) Heatmap representing the frequency that two introns within a read that spans at least four introns have the same splicing status (both spliced or both not spliced) in human K562 cells. Intron number represents the position of each intron within a read that spans at least four introns such that “intron 1” is the first intron and “intron 4” is the last intron within the set of four introns that a read spans.

## Discussion

nano-COP probes the relationship between transcription and splicing in living metazoan cells. This approach works by (1) combining two established approaches for purifying nascent RNA, 4sU labeling and cellular fractionation, thereby substantially improving the enrichment of unspliced pre-mRNA transcripts; and (2) sequencing the stringently purified nascent RNA directly through nanopores, which reveals both the location of transcription and the splice isoform of nascent RNA without amplification based biases. Importantly, treatment with the splicing inhibitor PlaB abolished evidence of co-transcriptional splicing detected by nano-COP, confirming that the approach effectively captures nascent RNA and is sensitive to changes in RNA processing rates.

nano-COP reveals the average physical proximity between nascent RNA ends (or Pol II position) and introns at the moment that splicing completes. We found that most introns in human cells were spliced after Pol II has transcribed 4 kb downstream, whereas in *Drosophila* S2 cells the majority of splicing occurred within 2 kb of introns. Given that the median transcription rate in both K562 and S2 cells is ∼1.25 kb per minute (Ardehali et al., 2009; O’Brien and Lis, 1993; Veloso et al., 2014), our splicing kinetics results are consistent with splicing half-lives estimated from metabolic labeling data: 2 and 14 minutes in *Drosophila* and mammalian cells, respectively (Pai et al., 2017; Rabani et al., 2014). Our findings suggest that in contrast to *S. cerevisiae* and *S. pombe,* in which splicing completes nearly immediately upon synthesis, in metazoan cells Pol II is not in close physical proximity to introns when splicing occurs (Carrillo Oesterreich et al., 2016; Herzel et al., 2018).

The differences observed by nano-COP between *Drosophila* and human cells are consistent with previously established models of intron and exon definition splicing, but provide novel insight in both cases (Figure 7). *Drosophila* has an abundance of short (<100 nt) introns that are assumed to be spliced through intron definition, where the spliceosome assembles at the 5’ and 3’SSs across the intron. Intron definition only requires the coordination of one intron at a time, so the reaction can be completed rapidly in the order of transcription, which we observe. We propose that the prominent role of cis-regulatory features (e.g. intron length and splice site sequences) in *Drosophila* splicing regulation is a consequence of intron definition–based splicing in which only one intron is recognized at a time. Human cells, on the other hand, have an abundance of long introns that are predominantly spliced through exon definition, in which the spliceosome initially assembles across exon boundaries before reorganizing to form cross-intron interactions and complete the splicing reaction. The greater distance between Pol II and introns at the time splicing occurs in human cells is in line with exon-definition splicing: the extra step of identifying exon boundaries before intron boundaries presumably increases the time required to splice each individual intron. Furthermore, the extra coordination of multiple introns during exon definition splicing likely relies on accessory factors to direct splicing across multiple introns. We propose that the simultaneous regulation of multiple introns by exon definition drives the coordinated splicing patterns that we observed across groups of three and four introns within human genes.

**Figure 7.**
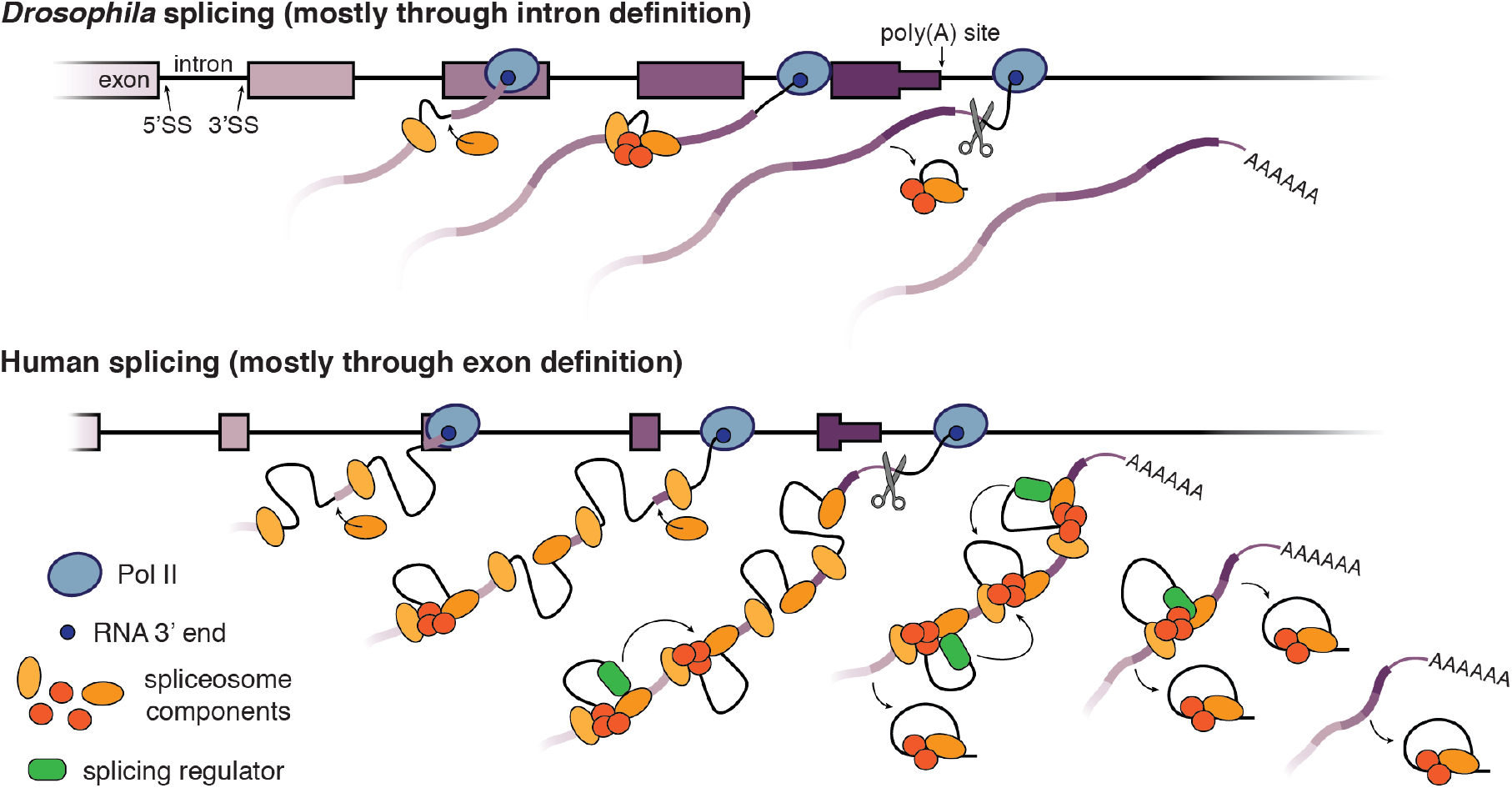
Coupling of transcription and RNA processing with intron definition versus exon definition splicing. A model depicting the differences in splicing regulation between *Drosophila* genes (top), which have an abundance of short introns that are spliced through intron definition, and human genes (bottom), which contain mostly long introns that are spliced through exon definition. In *Drosophila* cells, we observe that splicing typically occurs rapidly, co-transcriptionally, and in the order of transcription. By contrast, in human cells, splicing is slower such that intron splicing does not follow the order of transcription and terminal introns are spliced after cleavage and polyadenylation. Even though splicing catalysis is slower with exon definition, we propose that splicing factors still assemble on the transcribing RNA and are likely what is driving the regulated and coordinated splicing order we observe in human cells.

nano-COP data provide a new perspective on the early RNA processing steps that regulate intron splicing. Even though the transcription machinery is not in close physical proximity when splicing occurs in human cells, components of the spliceosome could still assemble on nascent RNA soon after introns are transcribed, in a manner that depends on transcriptional kinetics (Bentley, 2014). To complete intron splicing, the spliceosome likely relies on accessory factors such as RBPs that direct splicing to occur in a defined order within genes. We propose that both the rate of transcription and the order of intron splicing can regulate alternative splicing patterns and may contribute to may contribute to the tendency for introns neighboring alternative exons to experience splicing delays. As nanopore sequencing improves in the coming years, nano-COP will only increase in resolution and depth as deeper coverage will allow for analyses of specific introns, longer read lengths will expose patterns across larger transcripts, and greater accuracy will provide allele-specific information.

## Acknowledgements

We thank members of the Churchman lab, F. Winston, W. Timp, R. Workman, M. Marin, B. Smalec, and A. Markham for helpful discussions and advice; E. McShane, J. Bridgers, R. Ietswaart, K. Tyssowski, and C. Patil for critical reading of the manuscript; G. Amador and the DRSC/TRiP facility for guidance culturing *Drosophila* S2 cells; G. Yeo and E. Wheeler for processed eCLIP data. This work was supported by the NIH (R21-HG009264 and R01-GM117333 to L.S.C.; F31-GM122133 to H.L.D.) and the Fonds de Recherche du Québec - Santé (Post-doctoral fellowship award to K.C.).

## Materials and Methods

### Cell culture

K562 cells (ATCC, CCL-243) were maintained at 37°C and 5% CO_2_ in RPMI 1640 medium (ThermoFisher, 11875119) containing 10% FBS (ThermoFisher, 10437036), 100 U/ml penicillin and 100 ug/ml streptomycin (ThermoFisher, 15140122). S2 cells (Expression Systems, 94-005) were maintained at 25°C in Schneider’s *Drosophila* medium (ThermoFisher, 21720024) containing 10% heat inactivated FBS (ThermoFisher, 16140063), 50 U/ml penicillin and 50 ug/ml streptomycin (ThermoFisher, 15070063). To monitor co-transcriptional splicing in the presence of a splicing inhibitor, cells were incubated with 100 nM Pladienolide B (PlaB, Santa Cruz Biotechnology, sc-391691) or 0.01% DMSO (Sigma, D2438) for 1 hour, unless indicated otherwise.

### 4sU labeling

Cells were labeled in media containing 500 μM 4-thiouridine (4sU, Sigma, T4509) for 8 minutes. K562 cells were harvested in suspension at 0.8-1 million cells/mL. *Drosophila* S2 cells were labeled as an adherent layer at 95% confluency and lifted with gentle pipetting before harvesting. Cells were collected by centrifugation at 500 g for 2 minutes and washed once in 1X PBS. Samples for total or 4sU labeled RNA were immediately resuspended in Qiazol lysis reagent (Qiagen, 79306) for RNA extraction. Samples for chromatin-associated RNA purification proceeded immediately to the cellular fractionation protocol.

### Cellular fractionation

The cellular fractionation protocol was performed exactly as described in steps 8-21 of (Mayer and Churchman, 2016). In brief, samples with 10 million K562 cells or 50 million S2 cells were lysed for 2 min with 200 μl cytoplasmic lysis buffer (0.15% (vol/vol) NP-40 (Thermo Fisher Scientific, 28324), 10 mM Tris-HCl (pH 7.0), and 150 mM NaCl), layered over 500 μl of a sucrose cushion (10 mM Tris-HCl (pH 7.0), 150 mM NaCl, 25% (wt/vol) sucrose), and nuclei were collected by centrifugation at 16,000 g for 10 minutes. The nuclei pellet was resuspended in 800 μl wash buffer (0.1% (vol/vol) Triton X-100, 1 mM EDTA, in 1X PBS) and collected by centrifugation at 1,150 g for 1 minute. Washed nuclei were resuspended in 200 μl glycerol buffer (20 mM Tris-HCl (pH 8.0), 75 mM NaCl, 0.5 mM EDTA, 50% (vol/vol) glycerol, 0.85 mM DTT), and mixed with 200 μl nuclei lysis buffer (1% (vol/vol) NP-40, 20 mM HEPES (pH 7.5), 300 mM NaCl, 1 M urea, 0.2 mM EDTA, 1 mM DTT) before pulse vortex and incubation on ice for 2 minutes. The chromatin pellet was collected by centrifugation at 18,500 g for 2 minutes and resuspended in 1X PBS. All steps were performed at 4°C and all buffers were prepared with 25 μM α-amanitin (Sigma, A2263), 0.05U/μl SUPERase.In (Thermo Fisher Scientific, AM2694) and protease inhibitor mix (Roche, 11873580001).

### Western blot

Samples for western blot analysis were mixed with 1X PBS to equal volumes for all fractions. The chromatin sample was homogenized by adding 2 μl Benzonase (Sigma, E1014) and mixing with a 1-ml syringe and 22G needle. Western blots were performed using an antibody against Ser2-phosphorylated CTD (transcribing Pol II) (Active Motif, 3E10) or GAPDH (LifeTechnologies, AM4300/6C5) and imaged using a Licor Odyssey.

### Splicing inhibitor

*RT-PCR.* K562 and S2 cells were incubated with 0.1% DMSO or 10 nM, 100 nM, or 1 μM PlaB splicing inhibitor for 1 hour. The cells were subjected to cellular fractionation to collect cytoplasmic and chromatin-associated RNA. Both RNA samples were extracted using Qiazol lysis reagent (Qiagen, 79306) following the manufacturer’s instructions and DNA was degraded with DNase I (Qiagen, 79254). RNA was reverse transcribed with the SuperScript III First-Strand Synthesis System (Invitrogen, 18080051) using 5 ng/ul random hexamer primers following manufacturer’s instructions. PCR was performed with Phusion High Fidelity polymerase (ThermoFisher, 5F30S), 0.5 μM primers, and 1 μl cDNA from the RT reaction for 1 cycle of 2 min 98°C, 30 cycles of 20 sec 98°C, 30 sec 60°C, and 1.5 min 72°C and 1 cycle of 3 min 72°C. Samples were run on a 1% agarose gel to separate DNA fragments. Primer sequences include: BRD2_F CAAAATTATAAAACAGCCTATGGACATG and BRD2_R TTTTCCAGCGTTTGTGCCATTAGGA; Set_F AAATTGATGCCTGCCAGAAC and Set_R GGATTCTCGTCGAAATGGAA.

### Biotinylation and labeled RNA selection

RNA was first extracted using Qiazol lysis reagent (Qiagen, 79306) following the manufacturer’s instructions. Total RNA (∼300 μg per reaction) and chromatin-associated RNA (∼50-60 μg per reaction) were subjected to 4sU labeled RNA purification as previously described (Dölken et al., 2008; Schwalb et al., 2016). In brief, labeled RNA (1 μg / 10 μl) was incubated with 10% biotinylation buffer (100mM Tris pH 7.5, 10mM EDTA) and 20% EZ-Link Biotin-HPDP (1 mg/mL resuspended in DMF, Thermo Fisher Scientific, 21341) for 1.5 hours at 800 rpm and 24°C in the dark. RNA was purified by mixing with a 1:1 ratio by volume of chloroform/isoamylacohol (24:1), separating with a phase-lock tube at 16,000 g for 5 min, and performing isopropanol precipitation. Biotinylated RNA separation was performed using the μMACS streptavidin kit (Miltenyi Biotec, 130-074-101). RNA was mixed with μMACS streptavidin beads at a 2:1 ratio by volume at 800 rpm and 24°C for 15 min. RNA-streptavidin beads mix was transferred to the μMACS column and washed with wash buffer (100 mM Tris pH 7.5, 10 mM EDTA, 1 M NaCl, 0.1% Tween 20) at 65°C and room temperature each 3 times. Selected RNA was eluted off the magnet with 0.1M DTT and purified using the miRNeasy micro kit (Qiagen, 217084) with on-column DNase I treatment (Qiagen, 79254).

### Illumina sequencing libraries preparation and sequencing

Libraries were prepared using the Ovation Universal RNA-seq System (NUGEN, 0343-32) with Universal Human rRNA strand selection reagent (NUGEN, S01859) following the manufacturer’s instructions. All samples were sequenced 2×80 on a NEXTseq 500 sequencer (Illumina, San Diego, CA, USA) in the Biopolymers Facility at Harvard Medical School.

### Illumina sequencing data analysis

Paired-end reads were aligned to ENSEMBLE GRCh38 (release-86) and FlyBase dm6 (r6.19) reference genomes using STAR (v2.5.1a) (Dobin et al., 2013) with default parameters (except for readFilesCommand=cat, limitIObufferSize=200000000, limitBAMsortRAM=64000000000, outReadsUnmapped=Fastx, outSAMtype=BAM SortedByCoordinate, outSAMattributes=All, outFilterMultimapNmax=101, outSJfilterOverhangMin=3 1 1 1, outSJfilterDistToOtherSJmin=0 0 0 0, alignIntronMin=11, alignEndsType=EndToEnd). Total RNA RPKM was measured by counting the number of read pairs within the coding region of each gene divided by the length (in kilobases) of the gene divided by million mapped reads for each sample. Global percent spliced calculations were determined by dividing the number of spliced reads by total (spliced + unspliced) reads that span 3’SS junctions. Splicing index calculations were determined for each gene by counting the number of read pairs that span exon junctions by at least 3 nucleotides and measuring the number of spliced reads divided by unspliced reads; splicing index = 2 × spliced read pairs / (5’SS unspliced + 3’SS unspliced read pairs). NP-seq and mNET-seq datasets from HeLa cells treated with 0.1% DMSO or 1 μM PlaB for 4 hours were retrieved from (Nojima et al., 2015) under the GEO accession number GSE60358. Illumina total RNA sequencing from K562 cells treated with 100 μM 4sU for 4 hours were retrieved from (Schofield et al., 2018) under the GEO accession number GSE95854.

### Oxford Nanopore direct RNA library preparation and sequencing

Ribosomal RNAs were depleted from the 4sU-selected chromatin-associated RNA sample using Ribo-Zero Gold rRNA Removal kit (Illumina, MRZG126) for unperturbed samples or RiboMinus Eukaryotic Kit v2 (ThermoFisher, A15020) for PlaB/DMSO samples. Unless described otherwise, a poly(A) tail was added to the 3’ ends of the 4sU labeled chromatin-associated RNA with *E. coli* poly(A) polymerase from New England Biolabs (M0276S) for unperturbed samples or Clontech/Takara (2180) for PlaB/DMSO samples, incubating at 37°C for 1 hour or 7.5 minutes, respectively. The direct RNA sequencing protocol using the SQK-RNA001 kit (Oxford Nanopore Technologies Ltd.) for unperturbed samples or SQK-RNA002 kit (Oxford Nanopore Technologies Ltd.) for PlaB/DMSO samples was followed exactly as described by the manufacturer with minor modifications described below. In brief, 500 ng of RNA sample was ligated to the provided splinted adapter with T4 DNA ligase for 15 minutes at room temperature. Ligated RNA sample was reverse transcribed using SuperScript III (Invitrogen, 18080044), as recommended by Oxford Nanopore Technologies to improve the reading of RNA through the nanopore. Samples were purified with Agencourt RNAClean XP beads (Beckman Coulter, A63987) and ligated to the sequencing adapter with preloaded motor protein. After a second purification step with Agencourt RNAClean XP beads, the sample was resuspended in Elution Buffer, mixed with RNA Running Buffer, loaded onto a primed FLO-MIN106 flowcell, and sequenced using MINKNOW software for 48 hours with default settings for direct RNA sequencing. nano-COP datasets from human K562 cells and *Drosophila* S2 cells were prepared in biological triplicates. An additional sample from *Drosophila* S2 cells was prepared as a technical replicate and was combined with the other technical replicate for downstream analyses. nano-COP datasets from each cell type treated with 0.01% DMSO and 100 nM PlaB were prepared in biological duplicates.

### Direct RNA nanopore sequencing basecalling and alignment

Raw signal fast5 files from the unperturbed samples were basecalled using Albacore 2.2.7 (Oxford Nanopore Technologies Ltd.) with the following parameters: read_fast5_basecaller.py --flowcell FLO-MIN106 --kit SQK-RNA001 --recursive --output_format fast5,fastq --worker_threads 8 --save_path ${savePath} --input ${inputPath}. PlaB/DMSO sequences were collected by live basecalling with MinKNOW (release 18.12.6 and 18.12.9). To enable sequence alignment, RNA sequences that pass basecalling thresholds were converted into DNA sequences by substituting U to T bases before mapping. The reference genomes used for K562 and S2 sequence alignments were obtained from ENSEMBLE GRCh38 (release-86) and FlyBase dm6 (r6.19), respectively. Sequences were aligned to the reference genomes using minimap2 (version 2.10-r764-dirty) (Li, 2018) with recommended parameters for Oxford Nanopore Technologies direct RNA sequencing (-ax splice -uf -k14) and GMAP (version 2018-03-25) (Wu and Watanabe, 2005) with default parameters. We observed consistent results between the two aligners and decided to display all analyses using minimap2. All analyses were performed using reads that pass the MINKNOW sequencing threshold (QC>7) and align uniquely to the genome.

### Read coverage per gene visualization and comparisons

To display nano-COP reads for figures, split bed files with read information were converted into diagrams using pyGenomeTracks (Ramírez et al., 2018). For read coverage per gene comparisons, nano-COP datasets from replicates 1 and 3 were downsampled to have the same number of reads per run. All sequences from K562 nano-COP replicates were combined when comparing with Illumina sequencing data. Coverage per gene from ONT and Illumina samples was calculated using BEDTools coverage (Dale et al., 2011; Quinlan and Hall, 2010) with parameters s=True and mean=True.

### Comparisons between nano-COP and direct RNA sequencing of poly(A)-selected mRNA

Direct RNA sequencing samples with poly(A) selected RNA from immortalized human B-lymphocyte cells (GM12878) by the Oxford Nanopore RNA Consortium (Hopkins run 1 and UCSC run 1) (Workman et al., 2018) were accessed from https://s3.amazonaws.com/nanopore-human-wgs/rna/fastq/Hopkins_Run1_20170928_DirectRNA.pass.dedup.fastq and https://s3.amazonaws.com/nanopore-human-wgs/rna/fastq/UCSC_Run1_20170907_DirectRNA.pass.dedup.fastq. The reads were aligned to the hg38 reference genome using the same parameters for nano-COP as outlined above. Direct RNA reads from nano-COP and Oxford Nanopore RNA Consortium mRNA were analyzed for ‘match percent’, which represents the number of bases that match the reference genome divided by the total number of bases that align to the reference genome for each read. Samples were also compared for ‘read length’, as determined by the length of the read in the fastq file. To generate confusion matrices from each library to compare sequencing and alignment accuracies, all mapped regions of direct RNA nanopore reads were recorded for their aligned base and reference base. For 100,000 random aligned sequence segments in each sample, the frequency of each base matching the reference was recorded and plotted as a matrix.

### Direct RNA 3’ end analysis

RNA 3’ ends (deriving from the 5’ ends of sequenced reads) were recorded from minimap2 sequence alignments and assigned to gene feature categories. “Intron” and “exon” regions refer to their respective annotated features within protein coding genes from hg38 RefSeq annotations. “Poly(A)” sites are defined as regions within 50 nucleotides of the end coordinate of annotated genes. In K562 datasets, the poly(A) region also includes coordinates within 50 nucleotides of RNA-PET annotations from cytoplasm and chromatin fractions in K562 ENCODE data (ENCODE Project Consortium, 2012). “Post-poly(A)” sites are defined as the region between 50-550 nucleotides after the end of annotated genes. “Splice sites” are defined as 50 nucleotides upstream and 10 nucleotides downstream of annotated 5’ splice sites. “Undetermined” categorizes reads that align to more than one category and “other” represents read ends that do not align in the sense direction of annotated gene features (e.g. antisense transcripts, noncoding RNAs, intergenic transcription, etc.). Poly(A) tail lengths were estimated using Nanopolish version 0.11.1 (Loman et al., 2015; Workman et al., 2018). Raw signal fast5 files were indexed with *nanopolish index*; reads were segmented and poly(A) tail lengths were calculated with *nanopolish polya* using default parameters. Reads with the quality control flag “PASS” and with estimated tail lengths greater than 0 were used to plot tail length distributions for RNA 3’ ends aligning to gene bodies (intron, exon or splice site), poly(A) sites, or post-poly(A) sites based on annotated features from hg38 RefSeq annotations as described above. SIRV sequences and annotation files were downloaded from the Lexogen website (https://www.lexogen.com/wp-content/uploads/2017/06/SIRV_Set1_Sequences_170612a-ZIP.zip). Direct RNA nanopore sequencing data from SIRV E0 was taken from (Garalde et al., 2018) under the sra accession number SRR6058584. Reads were aligned using the same default parameters of minimap2 (Li, 2018) for nano-COP as outlined above. The 3’ end positions of reads were plotted in relation to the encoded poly(A) sites of three SIRV transcripts (SIRV101, SIRV504, and SIRV701).

### Splicing intermediates analysis

The prevalence of splicing intermediates, which are reads deriving from the splicing process rather than active transcription, were observed by measuring the coverage of read ends (5’ and 3’) at both splice sites and the middle of introns across the genome. RNA 3’ ends that align exactly at 5’SS (or the last base of an exon) are considered to be splicing intermediates because they likely arise from a free RNA end during the splicing reaction rather than active transcription. With the assumption that the downstream intron is actively undergoing splicing when the 3’ end of the read is at the 5’SS of the downstream intron, splicing order was determined for these intermediates by measuring the splicing status of the upstream (first transcribed) intron. Splicing status and order were implemented as described above.

### Identifying constitutively spliced introns for splicing analyses

Constitutively spliced introns were identified using total RNA sequencing data from human K562 cells in this study and *Drosophila* S2 cells from (Pai et al., 2017) under the GEO accession number GSE93763. RefSeq annotations of all gene and intron coordinates were extracted from the UCSC table browser both for hg38 and dm6. Constitutively spliced introns were determined to be those that have at least 20 reads that span either splice junction by at least 4 nt overlap and more than 80% of the spanning reads are spliced. These constitutively spliced introns were labeled as the “medium stringency” introns and used in all analyses for the distance transcribed before splicing plots. “High stringency” (5 nt overlap; 50 read coverage; 90% spliced), “low stringency” (3 nt overlap; 10 read coverage; 50% spliced), and “no stringency” (3 nt overlap; 0 read coverage, and 0% spliced) intron datasets were also produced to demonstrate the variation in results with different levels of intron retention. Maximum stringency categories for all introns are reported in Supplemental Tables 1 and 2.

### Determining splicing status of introns in Oxford Nanopore direct RNA sequencing reads

Due to the high error rate of nanopore sequencing, custom scripts were utilized to determine splicing status of direct RNA nanopore reads. First, aligned reads that overlap annotated intron coordinates (see above section for details on included introns) were identified and characterized using BEDTools intersect (Dale et al., 2011; Quinlan and Hall, 2010). Features of read cigar strings were extracted for the 50 nt around the intron 5’ and 3’ splice sites and the entirety of the intron using the Pysam toolkit (https://github.com/pysam-developers/pysam; (Li et al., 2009)). Reads were called as ‘not spliced’ if the alignment file shows no indication of splicing (CIGAR string “N” = 0) within the 50 nt around each splice site, mapped portions of the read (rather than deletions) represent greater than 50% of the 50 nt around each splice site, and at least 75% of the read within the intron is aligned to the reference. Reads were called as ‘spliced’ if the alignment file displays the start or end of a splicing event within the 50 nt around both splice sites and the size of the aligned splicing event is within 90-110% and 100 nt of the intron size. If aligned reads that map to introns do not meet these qualifications, the splicing event is characterized as ‘undetermined’ and not used in subsequent analyses.

### Measuring the distance transcribed before splicing

Reads with transcript 3’ ends that map near poly(A) sites or splice sites were removed from this analysis since it is likely they do not correspond to actively transcribing Pol II. Poly(A) sites were defined as the RefSeq annotated end coordinate of a gene from hg38 and dm6 genome assemblies as well as coordinates of RNA-PET 3’ end signal from cytoplasm and chromatin fractions in K562 ENCODE data (ENCODE Project Consortium, 2012). Reads with 3’ ends within 150 nt upstream or any distance downstream of an annotated poly(A) site were discarded from the analysis. To discard reads that possibly originated from splicing intermediates, RNA sequences with 3’ ends between 50 nt upstream and 10 nt downstream of an annotated intronic 5’ splice site were discarded from the analysis. Reads with more than 150 nt soft-clipped from the RNA 3’ ends during alignment were also discarded from the analysis due to uncertainty in Pol II position. RNA 3’ end mapping was performed with BEDTools intersect (Dale et al., 2011; Quinlan and Hall, 2010). The remaining reads were characterized for intron coverage and splicing status as described previously. In order to avoid read length constraints biasing the distance transcribed before splicing results, we only included cases where the length of the read is greater than the genetic distance from the read end to the intron 3’SS by at least 150 nt. Thus, the intron would be measured regardless of whether or not other introns within the same read are spliced or not spliced. For all reads included in the analysis, the distance between the 3’ end of the read and the 3’SS of the intron(s) it aligns to as well as the splicing status of the intron(s) it aligns to were recorded. Transcribed distances past intron 3’SS’s were binned into 100-500 nt windows depending on the number of datapoints in each bin. For all points within a bin, the percent of spliced introns was measured such that percent spliced = number of spliced molecules / total number of molecules.

### Characterizing the relationship between transcription termination and last intron splicing

For this analysis, poly(A) sites were defined as the end coordinate of RefSeq annotated genes from hg38 and dm6. Only the 3’-most poly(A) site for each terminal intron was utilized in this analysis. Reads with 3’ ends within 50 nt upstream and 50 nt downstream of poly(A) sites are referred to as “cleaved” and likely represent 3’ end processed and polyadenylated transcripts. Reads with 3’ ends between 50 to 500 nt downstream of poly(A) sites are referred to as “uncleaved” and likely represent pre-cleaved nascent transcripts. For reads in these two classes that also span the 3’SS of the terminal intron within the same gene, the splicing status of the intron it spans was determined as described above. The proportion of spliced terminal introns out of total was determined and plotted for each class and species.

### Determining alternatively spliced introns

Alternative exons were identified from total RNA sequencing data in K562 cells from this study and S2 cells from (Pai et al., 2017) using MISO (Katz et al., 2010). Mean insert length and standard deviation were assessed using CollectInsertSizeMetrics from Picard v.18.15 (http://broadinstitute.github.io/picard). Alternative event annotations (skipped exon, alternative 5’ or 3’ splice sites, and mutually exclusive exons) were obtained from the MISO annotations webpage (https://miso.readthedocs.io/en/fastmiso/annotation.html) for dm3 and hg19 (version 2) and lifted over to dm6 and hg38 using CrossMap (Zhao et al., 2014). Only alternative exons with percent spliced in (PSI) > 0.8 in the corresponding cell type (K562 or S2 cells) were considered for further analyses in order to exclude the confounding effect of partial splicing on kinetics measurements. In addition, annotations of all gene, transcript and exon coordinates were obtained in GTF format from NCBI (RefSeq) for hg38 and from FlyBase for dm6. For S2 cells, introns were considered alternative when they adjoin exons that are not present in every transcript of an annotated RefSeq gene and are identified as alternative by MISO with PSI > 0.8. For alternative 5’ or 3’ splice sites, only introns in 3’ or in 5’ of the alternative exon, respectively, were considered alternative. Conversely, introns were considered constitutive when they adjoin exons that are present in every isoform of an annotated RefSeq gene and are not identified as alternative by MISO. Introns that did not meet these criteria were excluded from the analysis. For K562 cells, since most introns neighbor exons that are not present in every isoform of a gene, introns were only considered alternative when they adjoin exons that are identified as alternative by MISO with PSI > 0.8 and introns were considered constitutive when they adjoin exons that are not identified as alternative by MISO. Introns that did not meet these criteria were excluded from the analysis. For measuring the distance transcribed before splicing of reads spanning alternative and constitutive introns, we used the same approach described above. Alternative splicing classifications for all introns are reported in Supplemental Tables 1 and 2.

### Determining the order of intron splicing

Reads that span two or more introns were used to characterize the order of intron splicing by nanopore sequencing. The algorithm requires that the length of the read must be greater than the genetic distance (containing the intron sequences) between the 3’ end of the read and the 5’ splice site of the first intron transcribed in the pair. This step ensures that the size of either intron in the pair does not bias the order of splicing findings. For reads that span two or more introns, the splicing status of introns in each neighboring pair was recorded as described above. For neighboring introns that have different splicing statuses, the frequency at which the upstream intron in the pair (first transcribed) is spliced before the downstream intron in the pair (second transcribed) was recorded for each species.

### Intron features that influence splicing order and kinetics

RefSeq annotations of gene and intron coordinates were extracted from the UCSC table browser for hg38 and dm6. Features of all introns in K562 and S2 cells, such as lengths of introns and neighboring exons, were recorded to determine which cis elements influence splicing kinetics and/or order. 5’SS and 3’SS consensus sequence scores were extracted from hg38 and dm6 assemblies using MaxEnt with default parameters (Yeo and Burge, 2004). Intron position within genes (e.g. first, middle, and last) was also determined from the annotation file. Intron features are reported in Supplemental Tables 1 and 2.

### Analysis of eCLIP data in K562 cells

For assessing RNA-binding protein (RBP) density in K562 cells, the fold change over input for significant peaks in previously published enhanced crosslinking and immunoprecipitation (eCLIP) data was obtained for 120 RBPs (Van Nostrand et al., 2018). eCLIP peaks located in introns were identified using BedTools intersect, requiring that at least 50% of the peak overlap with the intron. For each intron, we calculated the sum of the density of all peaks within this intron for all RBPs. RBP density was set to 0 for introns that do not overlap any peaks. Total RBP density per intron is reported in Supplemental Table 1. To compare RBP density in pairs of spliced and unspliced introns, we used all reads that span two or more introns as described above. For each individual RBP, if at least two intron pairs displayed binding in one intron (n=93 RBPs), the difference in binding density between spliced and unspliced introns within pairs was assessed using a paired t-test. In addition, for RBPs binding at least one spliced and one unspliced intron (n=63 RBPs), the number of bound spliced and unspliced introns was compared using a chi-square test. Multiple testing correction was performed using the Bonferroni method.

### Random forest model to determine features that influence splicing order in pairs

The predictive value of intron features for splicing order was determined using a Random Forest model with python scikit learn (Pedregosa et al., 2011). Features for intron length, surrounding exon lengths, splice site scores, intron position within genes, and RBP density (K562 cells only) were compiled for all introns. Feature importance scores were generated using a random forest classifier with 75% training and 25% testing sets. The total reads spanning intron pairs in the model are 4,316 for S2 cells and 2,305 for K562 cells. To avoid the same intron pairs in training and test sets, the intron pairs were sorted by gene name with the parameter “shuffle=False” during splitting. Default parameters were used in the random forest classifier (except for random_state = None, n_estimators = 300, max_features = ‘log2’, max_depth = None). ROC curve and AUC measurements were determined from binary prediction probabilities. Prediction accuracy was determined by measuring the difference between the model’s predictions and measured values. The baseline score was determined using a “null” parameter that has the same value for every training and testing pair; thus, baseline represents the prediction accuracy with no additional information added to the model.

### Splicing coordination across multiple introns

Reads that span 3 or more introns were collected in the same manner as reads spanning intron pairs (described above) and used to characterize the coordination of intron splicing. In cases where at least one of the three introns in the triplet has a different splicing status than the others, the order of splicing was determined as described for intron pairs. The frequency of triplet and quadruplet splicing patterns was recorded and mapped as a bar plot for all samples in K562 and S2 cells. The heatmap of intron splicing comparisons was prepared by compiling all reads that span four introns where at least one of the four introns has a different splicing status than the others. In every read the splicing status of each intron was compared to the splicing status of each other intron within the same read. The frequency at which two distinct introns within a quadruplet have the same splicing status (e.g. both not spliced or both spliced) was plotted as a heatmap.

---

**Figure S1.**
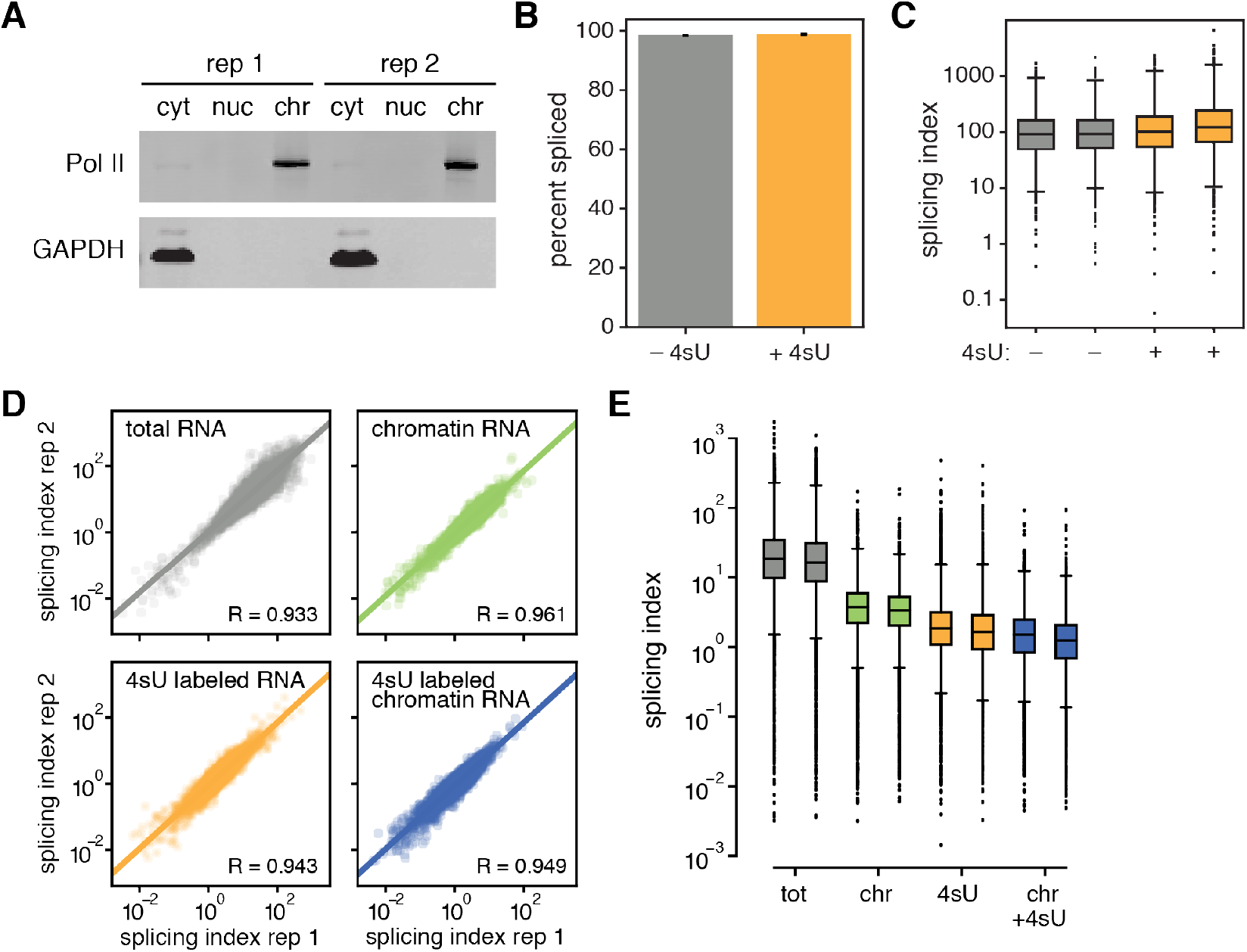
Nascent RNA enrichment with cellular fractionation and 4sU metabolic labeling (related to Figure 1). (A) Western blot of subcellular fractions from K562 cells with primary antibodies against Pol II Ser2 phosphoisoform (elongating Pol II) and GAPDH. cyt = cytoplasm; nuc = nucleoplasm; chr = chromatin. (B-C) Total RNA-seq datasets of cells incubated with and without 100 μM 4sU for 4 hours acquired from (Schofield et al., 2018) were compared for (B) global percent of spliced molecules and (C) splicing index. (D) Correlation plots of splicing index for each Illumina sequencing library. Pearson’s R is labeled at the bottom right of each plot. (E) Distribution of the splicing index for each sample across two biological replicates. tot = total RNA; chr = chromatin-associated RNA; 4sU = 4sU labeled RNA; chr+4sU = 4sU labeled chromatin-associated RNA. Percent spliced = spliced reads / total reads aligning to 3’SS junctions; splicing index = 2x spliced reads / unspliced reads that align to 5’ and 3’ splice junctions.

**Figure S2.**
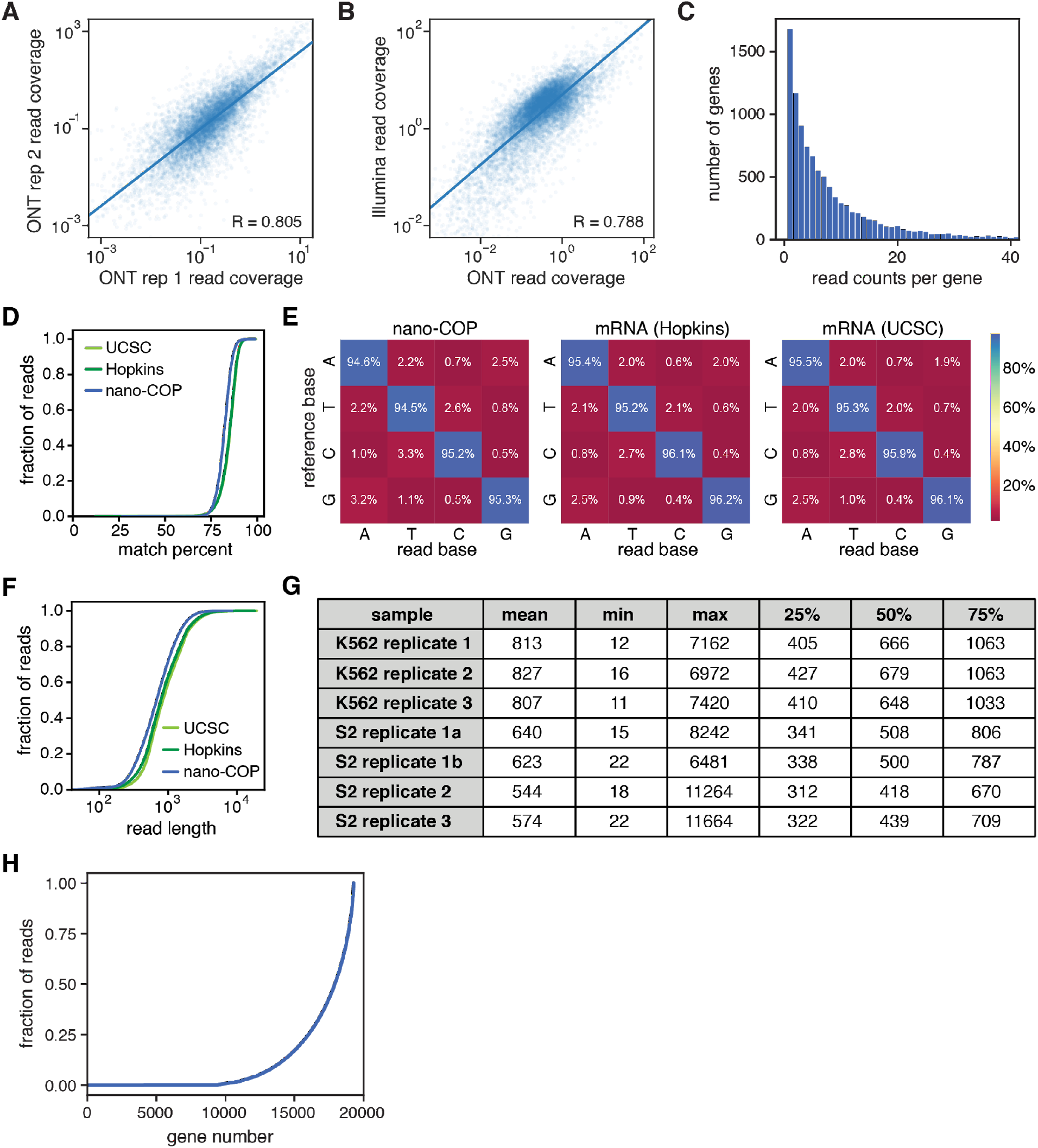
Key sequencing and alignment statistics of nano-COP (related to Figure 1). (A) Correlation plot of two nano-COP biological replicates from K562 cells by read coverage per gene; pearson’s R = 0.805. (B) Correlation plot of K562 4sU labeled chromatin-associated RNA sequenced on an Illumina instrument and Oxford Nanopore Technologies (ONT) MinION as read coverage per gene; pearson’s R = 0.788. (C) Histogram of nano-COP read counts per gene from replicate 1 in human K562 cells. (D) Cumulative distribution plot of alignment match percent (the percent of the read that exactly matches the reference sequence) from nano-COP and poly(A)-selected mRNA datasets. (E) Confusion matrix of read base calls versus reference bases for nano-COP and poly(A)-selected mRNA. (F) Cumulative distribution plot of direct RNA read lengths passing the default basecalling threshold from nano-COP and poly(A)-selected mRNA datasets. (G) Descriptions of ONT read lengths for all reads that pass the default basecalling threshold from human K562 and *Drosophila* S2 nano-COP datasets. Replicates 1a and 1b from *Drosophila* S2 cells were prepared as technical replicates. (H) Cumulative fraction of total reads that align to individual genes from nano-COP replicate 1 in human K562 cells. The direct RNA sequencing datasets of poly(A)-selected mRNA in D-F were generated as part of the Oxford Nanopore RNA consortium and originated from labs at the University of Southern California (UCSC) and Johns Hopkins (Hopkins) (Workman et al., 2018).

**Figure S3.**
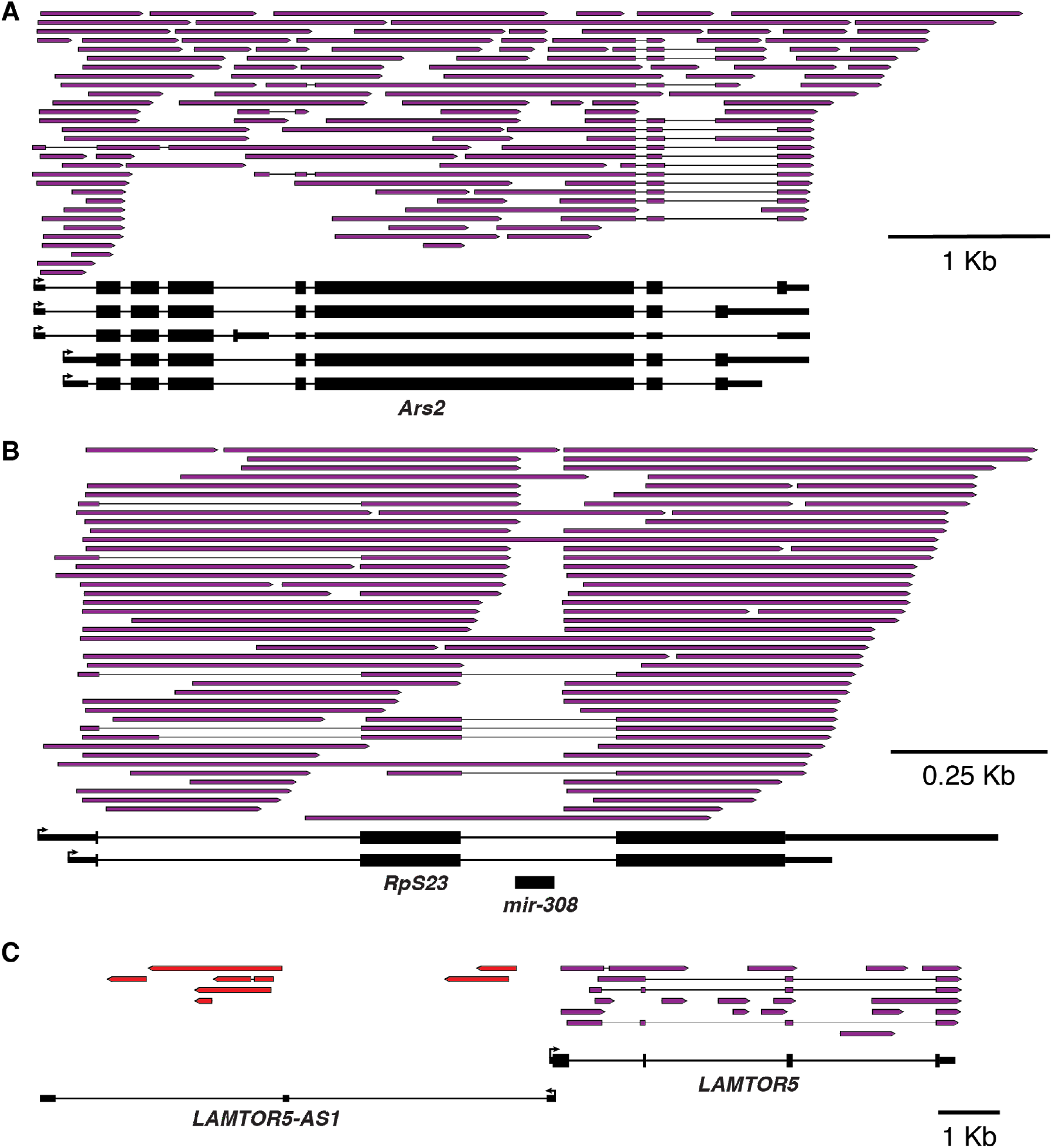
Example nano-COP reads reveal detection of alternative splicing, miRNA processing, and noncoding transcription (related to Figure 1). (A) Reads aligning to the *Ars2* gene in *Drosophila* S2 cells show alternative nascent isoforms in fruit flies. (B) Reads aligning to the *RpS23* gene in *Drosophila* S2 cells display processing around the *mir-308* miRNA. A pileup of read ends and starts around miRNA sites suggest a mechanism where pre-miRNAs are excised from intron segments before splicing completes. (C) Reads aligning to the *LAMTOR5* gene and *LAMTOR5-AS1* lncRNA in human K562 cells display nascent noncoding RNAs stemming from antisense transcription. Boxes represent read coverage and black lines represent skipped coverage due to splicing. Arrows on boxes represent the direction of transcription; purple boxes are reads transcribed in the sense direction of the coding gene; red boxes are reads transcribed in the antisense direction of the coding gene.

**Figure S4.**
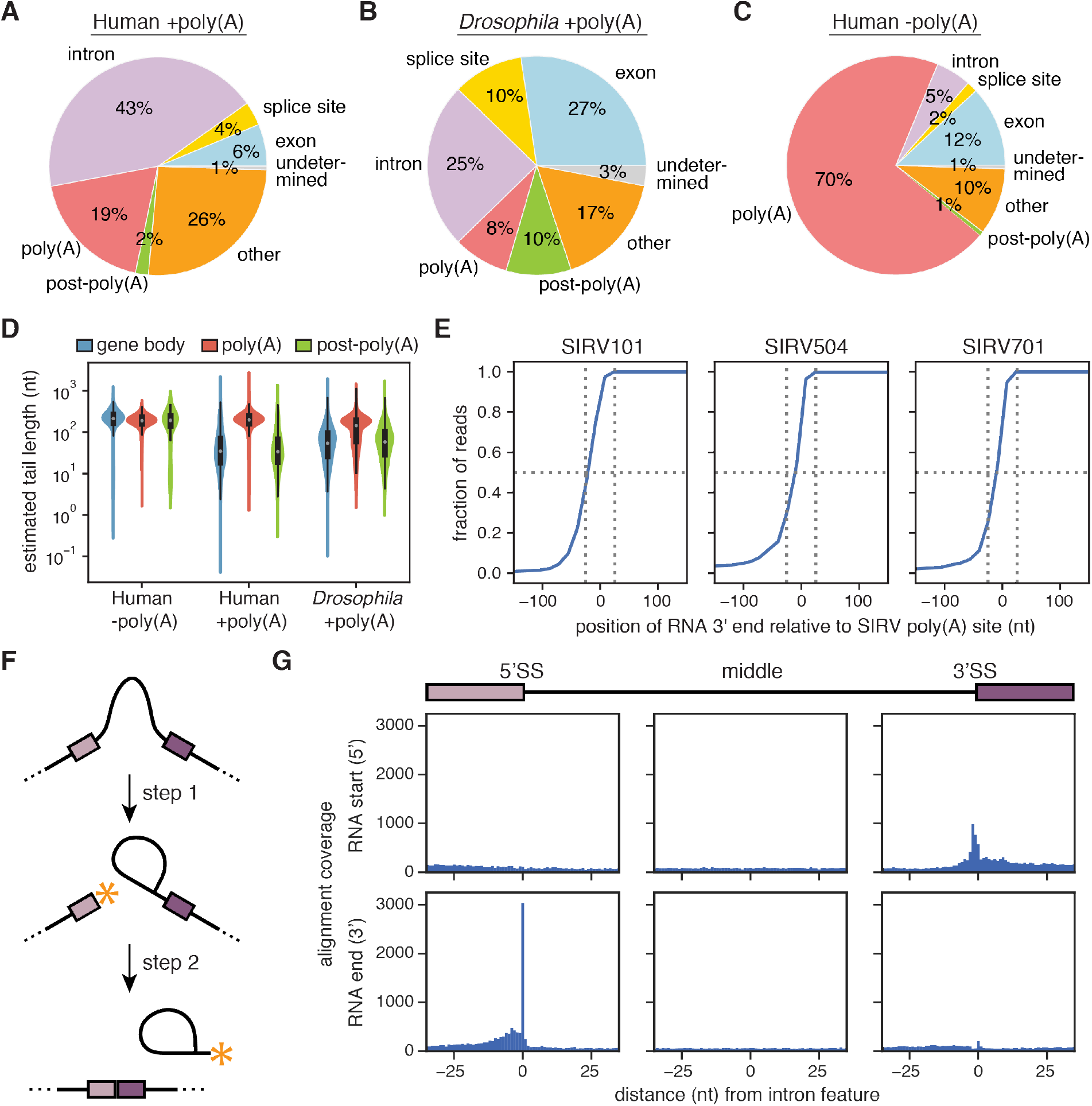
Alignment of nascent transcript 3’ ends in nano-COP data (related to Figure 2). (A–C) Distribution of nano-COP 3’ ends by nanopore sequencing in (A) human K562 cells with enzymatic poly(A) tail addition, (B) *Drosophila* S2 cells with enzymatic poly(A) tail addition, and (C) human K562 cells in the absence of enzymatic poly(A) tail addition. See Methods for descriptions of 3’ end alignment categories. (D) The length of poly(A) tails for sequenced RNAs with and without enzymatic poly(A) tail addition was estimated using nanopolish-polyA (Loman et al., 2015; Workman et al., 2018). Estimated tail lengths were plotted for RNAs in each sample that have 3’ ends aligning within gene bodies (exon, intron, or splice site), at poly(A) sites, or just downstream of poly(A) sites. (E) Cumulative distribution plot of transcript 3’ end positions from direct RNA nanopore sequencing of Spike-in RNA Variant (SIRV) control mixes aligned to the encoded poly(A) sites of three SIRV transcripts. Vertical dotted lines are at −25 and 25 nt from annotated poly(A) site; horizontal dotted line is at 0.50. (F) Cartoon demonstrating the catalytic steps of RNA splicing. Asterisks (*) represent available RNA ends during the splicing reaction. (G) The frequency of RNA starts (the 5’ end of the RNA when aligned) and ends (the 3’ end of the RNA when aligned) around intronic 5’ and 3’ splice sites and intron middles represented as histograms in human K562 cells. An increase in nascent RNA 5’ ends at 3’SS’s likely represents an interruption of the intron lariat within the nanopore. High coverage of nascent RNA 3’ ends at 5’SS’s likely result from free exon ends between the first and second catalytic steps of the splicing reaction.

**Figure S5.**
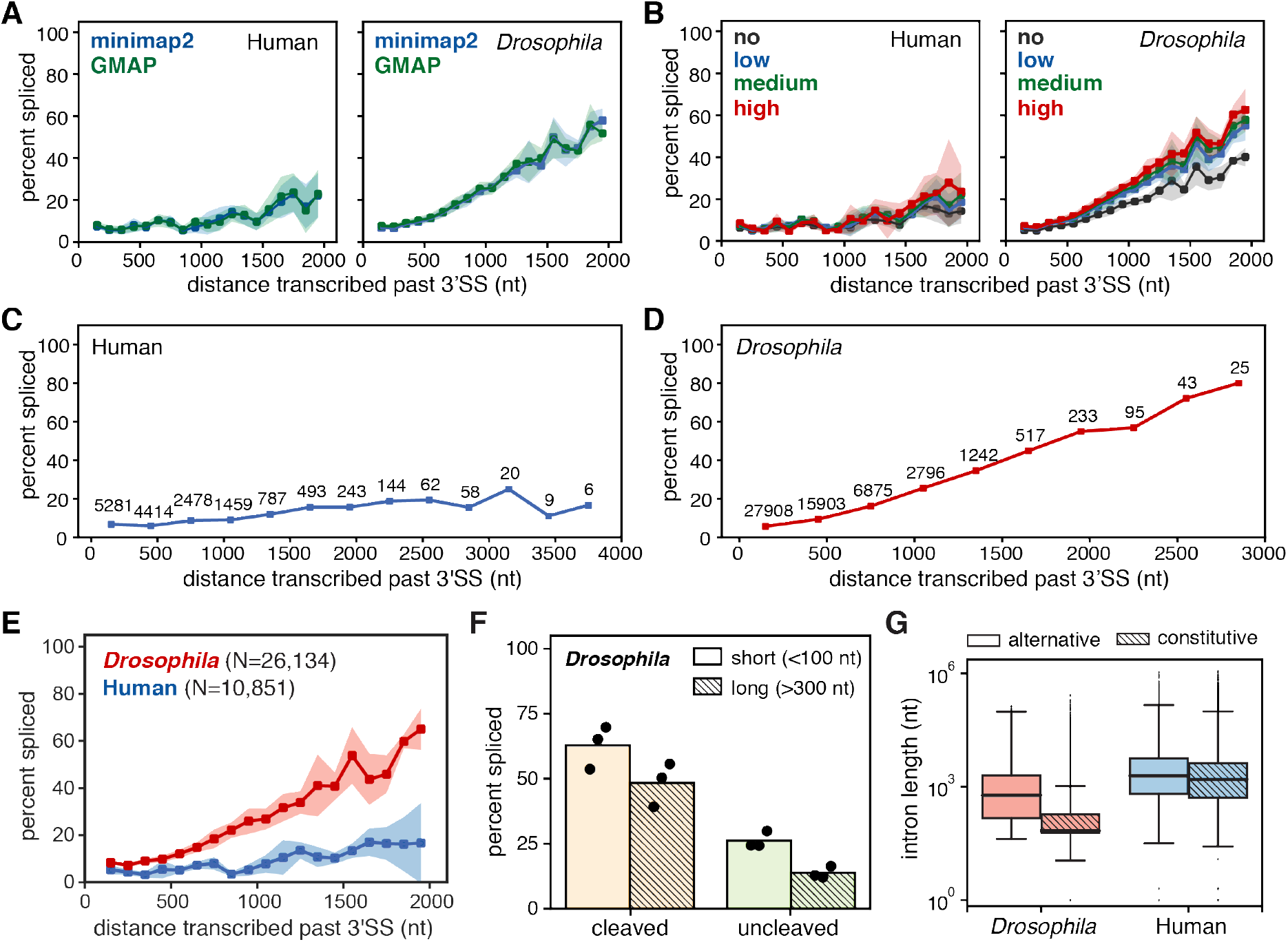
Extensions of plots that reveal the distance transcribed before splicing (related to Figure 2). (A) Global analysis of transcribed distance from 3’SS and the percent of spliced molecules with two different alignment programs, GMAP (Wu and Watanabe, 2005) and minimap2 (Li, 2018). (B) The distance transcribed before splicing with varying stringency levels of included introns. “No” stringency signifies that all introns were included. “Low”, “medium”, and “high” stringency levels include varying numbers of introns in the analysis based on intron retention levels from total RNA-seq data (see the Methods section for stringency descriptions). (C, D) Combining all biological and technical replicates to plot the global distance transcribed by percent spliced up to (C) 4 kb in human K562 cells and (D) 3 kb in *Drosophila* S2 cells. Numbers above points represent the total counts used to calculate the percent spliced measurement at each position. (E) The distance transcribed before splicing middle introns within genes after computationally removing first and last introns from the analysis. (F) The proportion of reads from *Drosophila* libraries with spliced terminal introns and 3’ ends in the “cleaved” bin (between 50 nt upstream and 50 nt downstream of poly(A) sites) and “uncleaved” bin (between 50-550 nt downstream of poly(A) sites). Bars are separated by intron length such that short introns (<100 nt) are represented as solid boxes while long introns (>300 nt) are represented as hashed boxes. Black points distinguish the results from three biological replicates. (G) Boxplots describe the lengths of introns that neighbor alternative versus constitutive exons in *Drosophila* (red) and human (blue) cells (t-test p-value < 10^−40^ in both species). Intron lengths are represented in log scale. Introns characterized as alternative are distinguished as empty boxes while constitutive introns are distinguished with hashed boxes.

**Figure S6.**
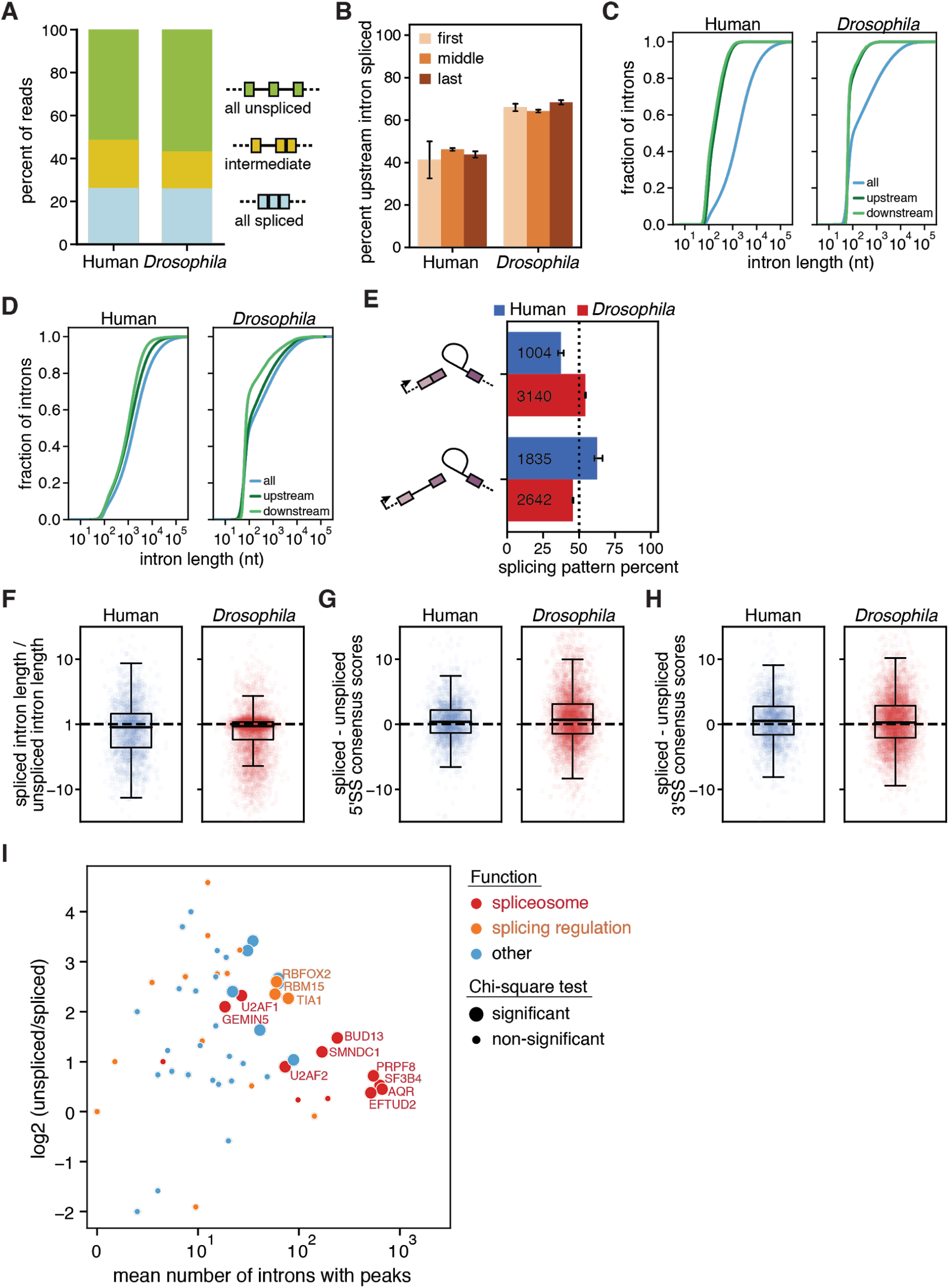
The features and regulators of splicing order (related to Figures 4 and 5). (A) Distribution of splicing patterns in nano-COP reads spanning at least two introns. “all unspliced” represents reads where every intron is present and therefore not spliced; “intermediate” represents reads where at least one intron is spliced and one intron is not spliced; “all spliced” represents reads where every intron within the read is spliced. (B) Bar plots to show changes in the order of splicing across intron pairs by position within the gene. “First” includes intron pairs where the upstream intron is the first intron within the gene; “middle” includes all intron pairs that do not contain a first or last intron within the gene; “last” includes all intron pairs where the downstream intron is the last intron within the gene. Error bars represent standard error of the mean across three biological replicates. (C) Cumulative distribution of intron lengths between all introns (blue) and only the upstream and downstream introns (green) analyzed within pairs. (D) Cumulative distribution of intron lengths between all introns (blue) and the upstream and downstream introns from RNAs that end at an intron 5’SS and are likely derived from splicing intermediates. The downstream intron is assumed to be in the process of splicing (between the first and second catalytic steps) while the upstream intron is either spliced or not spliced. (E) The frequency at which the upstream intron is spliced within reads deriving from splicing intermediates. (F) The percent change in intron length between first and second spliced introns within pairs. Human first spliced introns are shorter 55.1% of the time; *Drosophila* first spliced introns are shorter 61.1% of the time. (G, H) The difference in (E) 5’SS consensus scores and (F) 3’SS consensus scores between spliced and unspliced introns within pairs. In human cells, the spliced introns within pairs have higher 5’SS scores in 55.5% of cases and 3’SS scores in 54.8% of cases. In *Drosophila* cells, the spliced introns within pairs have higher 5’SS scores in 58.2% of cases and 3’SS scores in 53.0% of cases. (I) MA plot comparing individual RBP binding in spliced and unspliced introns. Each circle represents an RBP. The x-axis represents the mean number of spliced and unspliced introns with peak(s) while the y-axis represents the log2 ratio of the number of unspliced introns with peak(s) and spliced introns with peak(s). Larger circles show RBPs that bind more than five spliced and unspliced introns and that display statistically significant differences in a chi-square test comparing number of spliced and unspliced introns (Bonferroni-corrected p-value threshold = 0.00079). RBPs are colored based on their annotated functions in Table S1 from (Van Nostrand et al., 2018); functions that do not relate to “spliceosome” or “splicing regulation” are labeled as “other”.

**Figure S7.**
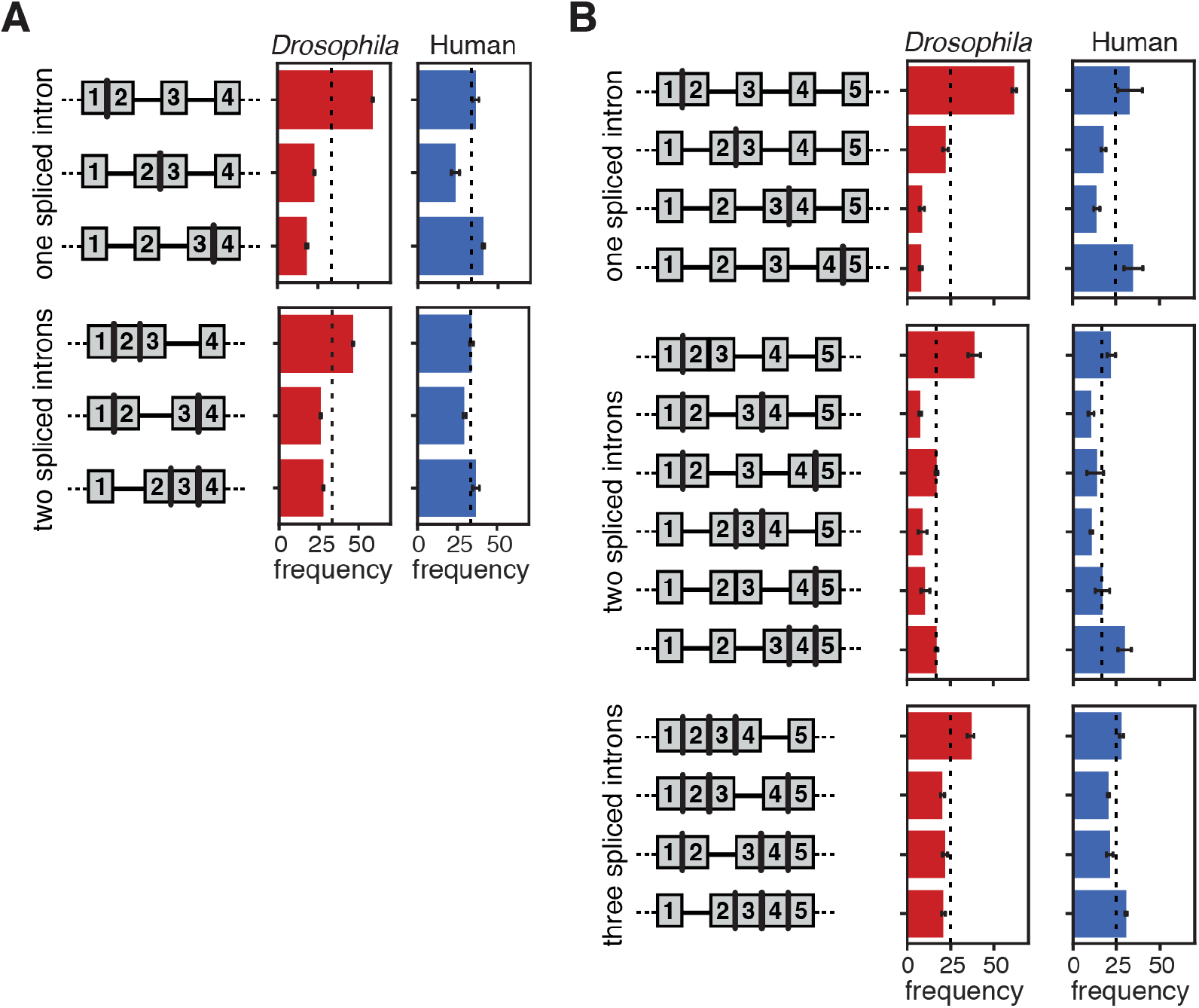
Characteristics of coordinated splicing across neighboring introns (related to Figure 6). (A–B) The proportion of reads in *Drosophila* S2 cells (red) and human K562 cells (blue) that exhibit the same splicing pattern as the schematic to the left across (A) intron triplets and (B) intron quadruplets when first and last introns within genes are not included in the analysis. Error bars represent standard error of the mean across three biological replicates.

## Supplementary Table legends

**Table S1. Features of all introns in human K562 cells.** The coordinates of the intron are displayed as chromosome, start position, end position, and strand. ‘Gene name’ refers to the RefSeq name for coding genes. ‘Intron pos’ is the intron number in the order of transcription. ‘Intron total’ is the total number of introns within that gene. ‘Intron order’ is the descriptive position of the intron within the gene (first, middle, or last). ‘Distance from start’ and ‘distance from end’ are the lengths in nucleotides from the start and end coordinates of the gene, respectively. ‘Intron length’, ‘upstream exon length’, and ‘downstream exon length’ are the sizes of the intron and surrounding exons in nucleotides. ‘5SS score’ and ‘3SS score’ are the splice site consensus sequence scores at the 5’SS and 3’SS, respectively, as determined by the maximum entropy calculations from (Yeo and Burge, 2004). RPKM refers to the reads per kilobase per million calculations from short read Illumina sequencing. ‘Splicing index’ refers to the splicing index results from short read Illumina sequencing. ‘Alt splicing’ signifies whether the intron neighbors an alternatively spliced exon in this cell line (refer to Materials & Methods). ‘Stringency’ refers to the lowest category of constitutive intron splicing stringency (as defined in the Materials & Methods section). ‘RBP density’ refers to the sum of fold change over input for all RBP peaks in the intron (from (Van Nostrand et al., 2018) and as defined in the Materials & Methods section).

**Table S2. Features of all introns in *Drosophila* S2 cells.** The coordinates of the intron are displayed as chromosome, start position, end position, and strand. ‘Gene name’ refers to the RefSeq name for coding genes. ‘Intron pos’ is the intron number in the order of transcription. ‘Intron total’ is the total number of introns within that gene. ‘Intron order’ is the descriptive position of the intron within the gene (first, middle, or last). ‘Distance from start’ and ‘distance from end’ are the lengths in nucleotides from the start and end coordinates of the gene, respectively. ‘Intron length’, ‘upstream exon length’, and ‘downstream exon length’ are the sizes of the intron and surrounding exons in nucleotides. ‘5SS score’ and ‘3SS score’ are the splice site consensus sequence scores at the 5’SS and 3’SS, respectively, as determined by the maximum entropy calculations from (Yeo and Burge, 2004). RPKM refers to the reads per kilobase per million calculations from short read Illumina sequencing from (Pai et al., 2017). ‘Splicing index’ refers to the splicing index results from short read Illumina sequencing. ‘Alt splicing’ signifies whether the intron neighbors an alternatively spliced exon in this cell line (refer to Materials & Methods). ‘Stringency’ refers to the lowest category of constitutive intron splicing stringency (as defined in the Materials & Methods section).

**Table S3. Characteristics of random forest models for splicing order.** Importance percentages for each intron feature represent the weight of each feature within the prediction model. For each feature, “1” represents the upstream intron (first transcribed) and “2” represents the downstream intron (second transcribed). The prediction accuracy for each random forest classifier from *Drosophila* and human intron pairs represents the percent of time the prediction is correct. Baseline represents the accuracy of predicting splicing order without any information from the feature inputs.

**Table S4. Comparison of RBP binding in pairs of spliced and unspliced introns within the same read.** RBPs that are significantly more likely to be bound to unspliced introns than spliced introns within pairs are shown. RBP binding was compared in two ways: 1) using a chi-square test comparing the number of spliced and unspliced introns bound by an RBP. Only RBPs with at least one bound spliced intron and one bound unspliced intron were considered (n=63); 2) using a paired t-test comparing RBP density in spliced and unspliced introns within the same pair. Only RBPs with binding in at least one intron of the pair in at least two pairs were considered (n=93). Multiple testing correction was performed using the Bonferroni method using alpha=0.05. The Bonferroni-corrected p-value thresholds for significance were 0.00079 (chi-square test) and 0.00054 (t-test). The same RBPs were statistically significant in both tests. The annotated function was obtained from Table S1 of (Van Nostrand et al., 2018) and annotations that do not relate to “spliceosome” or “splicing regulation” are labeled as “other”.

